# Integrated transcriptome-metabolome analysis reveals genotype-dependent temporal determinants of grapevine susceptibility to *Botrytis cinerea*

**DOI:** 10.64898/2026.11.26.766433

**Authors:** Clément Cuello, Benoit Monnereau, Samuel Bastos Serra Trinca, Cécile Gaillard, Vincent Lebeurre, Allegra Dovere, Bouziane Moumen, Claudia Rouveyrol, Pierre Pétriacq, Pierre Coutos-Thevenot, Sylvain La Camera

**Author notes:** Corresponding authors: Sylvain La Camera, Écologie et Biologie des Interactions, UMR7267 Université de Poitiers CNRS, F-86000 Poitiers, France. These authors contributed equally (Clément Cuello, Benoit Monnereau). UMR1236, SITI Laboratory, University Hospital, INSERM, EFS, Rennes, France.

## Abstract

Gray mold caused by *Botrytis cinerea* remains a major constraint for grapevine, yet the molecular basis of genotype-dependent susceptibility is poorly resolved. We investigated whether contrasting disease phenotypes are associated with pre-existing molecular states, differential deployment of shared responses, or genotype-specific defense programs. After screening seven *Vitis vinifera* cultivars and two American *Vitis* species, we selected *Vitis riparia*, *Vitis vinifera* ‘Merlot’ and ‘Syrah’ for comparative analysis of symptom development, fungal biomass, leaf transcriptomes and untargeted metabolomes before infection and at 24 and 48 h post-infection (hpi). *V. riparia* developed the smallest symptoms without reduced fungal biomass at 24 and 48 hpi, indicating a disease-tolerance component. Before infection, 5 341 genes were differentially expressed between *V. riparia* and both *V. vinifera* cultivars, and self-organizing maps revealed a distinct *V. riparia* transcriptional state that remained evident at 24 hpi. By 48 hpi, the transcriptional responses converged, with 9 426 differentially expressed genes shared among genotypes. Conserved late responses included repression of photosynthetic and chloroplast functions and coordinated activation of phenylpropanoid metabolism, lipid-derived signaling and cell-wall remodeling. Transcript-metabolite integration linked these programs to depletion of primary carbon metabolites, and accumulation of stilbenes and oxidized lipids. In parallel, *V. riparia* activated an additional defense module enriched in defense- and resistance-associated genes, detoxification and callose-related functions. These findings suggest that grapevine disease outcome is shaped by the interplay between basal molecular state and the kinetics of defense activation, rather than by the magnitude of a shared response.

**HIGHLIGHTS:**

- *Vitis riparia* limits necrosis despite similar *Botrytis* biomass
- Basal transcriptome and metabolome distinguish *V. riparia* before infection
- Late responses converge on chloroplast repression and defense metabolism
- Stilbenes, oxidized lipids and cell-wall programs define the shared response
- *V. riparia* activates an additional defense program potentially linked to tolerance

## 1. INTRODUCTION

Gray mold, caused by the necrotrophic fungus *Botrytis cinerea*, is one of the major diseases affecting grapevine, causing substantial losses in yield and fruit quality both before and after harvest (Williamson et al, 2007; Veloso and van Kan, 2018; Bi et al, 2023; Rahman et al, 2024). Susceptibility nevertheless varies considerably within the genus *Vitis*. Most cultivated *Vitis vinifera* genotypes are susceptible, although differences in disease severity occur among cultivars and can involve anatomical, chemical and physiological traits (Gabler et al, 2003; Pezet et al, 2003b; Rahman et al, 2020; Adrian et al, 2024). Wild *Vitis* germplasm displays a broader range of resistance phenotypes and represents an important resource for dissecting gray mold resistance. Screening of 41 *Vitis* genotypes identified several resistant Chinese wild accessions, whereas most tested *V. vinifera* genotypes or hybrids were susceptible (Wan et al, 2015). Consistently, *Vitis amurensis* has provided a useful model for identifying molecular mechanisms associated with resistance to *B. cinerea* (Wan et al, 2021). Resistance has also been described in Beta, an interspecific *Vitis riparia* x *Vitis labrusca* hybrid, in which two stable quantitative trait loci for gray mold resistance and associated candidate regulators were recently identified (Su et al, 2023). These studies illustrate the substantial genetic diversity available within *Vitis* for understanding the molecular determinants that shift the outcome of the interaction with *B. cinerea*.

Following inoculation, the grapevine-*B. cinerea* interaction progresses through a highly dynamic sequence of events, with an early 0-24 h post-infection (hpi) phase that is often largely asymptomatic, followed by increasing host-cell death and necrotrophic colonization (Veloso and van Kan, 2018; Bi et al, 2023; Rahman et al, 2024). Molecular studies in grapevine leaves, flowers and berries have revealed extensive reprogramming involving immune perception, hormone and redox signaling, cell-wall responses, lipid, carbon and specialized metabolisms, together with major changes in photosynthetic functions (Agudelo-Romero et al, 2015; Haile et al, 2017; Wan et al, 2021, 2024; Adrian et al, 2024; Rahman et al, 2024). Yet activation of canonical defense pathways does not necessarily result in reduced disease severity. The highly susceptible *V. vinifera* ‘Trincadeira’, for instance, undergoes extensive transcriptomic and metabolic reprogramming upon infection, including activation of phenylpropanoid metabolism and accumulation of trans-resveratrol and gallic acid, without preventing disease progression (Agudelo-Romero et al, 2015). Likewise, comparative studies indicate that both susceptible and resistant grapevines activate overlapping defense pathways, while differing in their timing, coordination and basal molecular state (Wan et al, 2021). Thus, disease outcome may depend less on whether a given defense pathway is activated than on when, to what extent and within which pre-existing molecular context it is deployed.

Comparisons of genotypes differing in susceptibility further suggest that the effectiveness of defense responses depends critically on their timing and underlying molecular context. In leaves, *V. amurensis* differs from susceptible *V. vinifera* before inoculation through basal transcriptional features associated with carbon and lipid metabolism, antioxidant capacity and immunity, and subsequently displays distinct early regulation of redox homeostasis, cell death and cell-wall defenses (Wan et al, 2021). Evidence for basal preconfiguration has also been obtained in fruit, although in a strong tissue- and developmental-stage-dependent context. In green berries, *V. vinifera* ‘Syrah’ displayed substantially fewer symptoms than *V. vinifera* ‘Trincadeira’ and this phenotype was associated with higher basal abundance or expression of components related to jasmonic acid/ethylene signaling, phenylpropanoid metabolism and putative defense metabolites (Soares et al, 2022). Earlier physiological comparisons similarly showed that comparable levels of latent *B. cinerea* infection can result in markedly different disease severity between grapevine cultivars, associated with differences in constitutive phenolic defenses and inhibition of fungal macerating enzymes (Pezet et al, 2003a). More recently, comparison of tolerant and susceptible *V. vinifera* genotypes showed that both could mount rapid early defense response to *B. cinerea*, whereas subsequent trajectories differed in the coordination of specialized metabolism, stress adaptation and growth-related processes (Maggiolini et al, 2025). Together, these studies indicate that susceptibility cannot be reduced to the intensity of a generic defense response. However, it remains unresolved whether contrasting disease phenotypes primarily reflect pre-existing molecular states, quantitative differences in shared responses, or activation of qualitatively distinct defense programs.

Moreover, integrated transcriptome-metabolome comparisons remain limited and have largely focused on reproductive tissues (Agudelo-Romero et al, 2015; Haile et al, 2020; Soares et al, 2022), leaving poorly resolved how transcriptional programs and metabolic changes are coordinated in leaves of genotypes displaying contrasting susceptibility. Recent applications of self-organizing map (SOM)-based transcriptome portrayal in grapevine have demonstrated its ability to resolve cultivar-specific and time-dependent transcriptional trajectories under environmental stress and across seasonal developmental transition (Konecy et al, 2023, 2024, 2026). However, to our knowledge, such an approach has not been used to resolve genotype-dependent transcriptional programs during the grapevine-*B. cinerea* interaction and directly associate them with paired metabolomic profiles. Here, we investigate how basal molecular configuration and subsequent transcriptional and metabolic trajectories relate to contrasting susceptibility to *B. cinerea*. To this end, we combined phenotypic characterization with transcriptomic and untargeted metabolomic profiling of leaves from *V. riparia* and *V. vinifera* ‘Merlot’ and ‘Syrah’ before infection and during early and late stages of the interaction. The underlying omics datasets are described in the associated data article (Monnereau et al, submitted to *Data in Brief*). Rather than considering differential-expression contrasts independently, we used SOM-based transcriptome portrayal to resolve coordinated basal, shared, and genotype-associated transcriptional trajectories, and directly associated module-level activity with metabolite accumulation profiles. This integrative framework allows us to distinguish basal, shared and genotype-specific transcriptional programs and to determine whether they are accompanied by coherent metabolic changes associated with contrasting disease phenotype.

## 2. MATERIAL AND METHODS

### 2.1. Plant and fungal materials

*V. vinifera* ‘Cabernet-Sauvignon’ grafted onto Fercal rootstock, ‘Chardonnay’, ‘Gamay’, ‘Merlot’, ‘Syrah’, ‘Cot Malbec’, and ‘Ugni Blanc’ grafted onto 3309 rootstock, together with non-grafted *V. riparia* ‘Gloire de Montpellier’ and *Vitis rupestris* ‘Rupestris du Lot’, were grown in 5.5-L pots containing a 3:1 (v/v) mixture of BP-Substrat 264 (Klasmann-Deilmann GmbH, Germany) and vermiculite, supplemented with 4 g.L-1 Osmocote exact standard 15-9-11 (12/14M) (Everris, Netherlands). Plants were maintained under greenhouse conditions with a 16-h light/8-h dark photoperiod and a minimum day/night temperature of 22C° and 18C°, respectively. *B. cinerea* strain B05.10 (Staats and van Kan, 2012) was recovered from conidial glycerol stocks and grown on Difco Potato Dextrose Agar (PDA) (Becton-Dickinson, United-States) at 22C° under long day conditions (16-h light/ 8-h dark) until sporulation. Conidia were harvested in sterile water and filtered through Miracloth (Merck, Germany).

### 2.2. Leaf discs infection for assessing susceptibility levels to *Botrytis cinerea*

Grapevine susceptibility to *B. cinerea* was assessed using the detached leaf-disc assay described in Monnereau et al (2026). Briefly, 2-cm-diameter leaf discs were excised from mature leaves, typically from nodes 8 to 10 from the shoot apex. They were placed, abaxial side up, on water-soaked Whatman 3 MM CHR paper, and inoculated with a 6-μl droplet of *B. cinerea* conidial suspension (5.10^5^ conidia.mL^-1^) prepared in 1:4 diluted Difco Potato Dextrose Broth (PDB; Becton-Dickinson, United-States). Inoculated discs were incubated for 72 h under water-saturated conditions. Necrotic areas were quantified by chlorophyll fluorescence imaging using an IMAGING-PAM M-series Chlorophyll Fluorometer (WALZ, Germany), as described previously (Monnereau et al, 2026). Briefly, discs were exposed to actinic light at 100 μmol.m^-2^.s^-1^ for 500 s, followed by a saturating pulse, and necrotic areas were quantified from [Y(NO)] images using ImageJ (Schneider et al., 2012) and the MicroArray Profile plugin.

### 2.3. Whole leaf infection and omics datasets

Whole-leaf infection assays used for omics analyses were performed as described in the associated data paper (Monnereau et al, *submitted to Data in Brief*) and following a previously published method from MetaboHUB-Bordeaux Metabolome (Hocini et al., 2026). Briefly, mature grapevine leaves, typically collected from nodes 8 to 10 from the shoot apex, were either immediately frozen in liquid nitrogen for the non-treated condition or inoculated on the abaxial side with *B. cinerea*. For infection assays, leaves were sprayed with a conidial suspension prepared at a concentration of 10^5^ in PDB1/4, whereas mock-treated leaves were sprayed with PDB1/4 without conidia. Leaves were incubated in water-saturated mini-greenhouses at 21°C and collected at 24 and 48 hpi. For each biological replicate, three leaves were pooled into a single sample and flash-frozen in liquid nitrogen.

RNA-seq library preparation, Illumina sequencing, read processing, mapping and gene-level quantification were performed as detailed in the associated data paper (Monnereau et al, *submitted to Data in Brief*). Briefly, strand-specific poly(A)-selected RNA-seq libraries were sequenced on Illumina NovaSeq 6000 platform in paired-end 2 x 150 bp mode. After quality control, trimming, filtering, and rRNA removal, reads were mapped to a concatenated reference genome comprising the *Vitis vinifera* ‘Pinot Noir’ T2T assembly and the *B. cinerea* B05.10 genome. Gene-level read counts were obtained using feature-based quantification, and the resulting count table was split into grapevine and fungal gene matrices for downstream analyses. *Vitis* gene-count matrix was retained and used for all downstream transcriptomic analysis. Raw RNA-seq data are available under BioProject accession PRJNA1404939.

Untargeted metabolomics profiling was performed on the same biological material, as described in the associated data paper (Monnereau et al, *submitted to Data in Brief*). Briefly, semi-polar metabolites were extracted from 10 mg of freeze-dried leaf powder with 80% ethanol containing 0.1% formic acid and methyl vanillate (250 µg.mL^-1^) as an internal standard. Extracts were filtered through a 0.22-µm membrane and analyzed by liquid chromatography coupled to high-resolution mass spectrometry in negative ionization mode using an Orbitrap Elite mass spectrometer. Chromatographic separation was performed on a Gemini C18 column (150 x 2 mm, 3 µm) using a water/acetonitrile gradient, and data-dependent MS/MS acquisition was performed at a resolution of 30 000 using stepped collision energies of 15, 30 and 45 eV. QC samples were injected every 10 samples to monitor analytical stability.

Raw LC-MS data were processed using MS-DIAL (v4.9.221218), and metabolomic features were filtered based on blank contribution, signal-to-noise ratio, and coefficient of variation across quality control samples. Before statistical analysis, feature intensities were corrected according to the initial dry mass for extraction, median-normalized, cube-root transformed, and Pareto-scaled. FDR-corrected ANOVA was performed in MetaboAnalyst (v6.0) to identify features showing significant variation across experimental conditions. Raw LC-MS data are available in MetaboLights under accession MTBLS16207.

### 2.4. *Botrytis cinerea* quantification in infected leaves

To quantify *B. cinerea* colonization in infected grapevine leaves, genomic DNA was extracted from non-treated grapevine leaves of each cultivar, *B. cinerea* mycelium, and infected leaf samples collected at 24 and 48 hpi. DNA extraction was performed using NucleoSpin Plant II kit (Macherey-Nagel, Germany), according to the manufacturer’s instructions. DNA concentration and quality were assessed by spectrophotometry using Multiskan GO µDROP system (Thermo Scientific, United Stated).

Quantitative PCR was performed on a LightCycler 480 thermocycler (Roche, Switzerland) using GoTaq qPCR Master Mix (Promega, USA). Organism-specific primers targeting *B. cinerea BcCutA* and grapevine *VvPDS* were used to quantify fungal and host genomic DNA, respectively. Primer sequences are listed in **Supplementary Table S1**.

Calibration curves were generated from DNA standards prepared using purified *B. cinerea* gDNA and gDNA extracted from non-treated grapevine leaves. These calibration curves were used to convert Ct values into DNA quantities in infected samples. For each infected sample, *B. cinerea* colonization was expressed as the ratio of fungal to grapevine gDNA, calculated as the *BcCutA*/*VvPDS* ratio. Three biological replicates were analyzed for each cultivar and timepoint. The significance of the genotype, sampling-time and genotype x time effects was assessed from the aligned rank transformed model. When appropriate, post-hoc genotype comparisons were performed separately at each time point with correction for multiple testing.

### 2.5. Differential expression analysis

Differential expression analysis was performed on the *Vitis* gene count matrix using DESeq2 (v.1.46.0, Love et al, 2014). Sample metadata were used to define a combined experimental group factor corresponding to each cultivar-by-condition combination. Genes with less than 10 counts in a minimum number of samples corresponding to the smallest experimental group size were filtered out. Differential expression was then tested using negative binomial generalized linear models implemented in DESeq2. Infection-responsive genes were identified independently for each cultivar and time point by contrasting infected samples with their corresponding mock-treated controls at 24 and 48 hpi. In non-treated conditions, pairwise comparisons were performed among the three grapevine genotypes, resulting in three genotype-by-genotype contrasts. Differentially expressed genes (DEGs) were defined by a Benjamini-Hochberg-adjusted p-value ≤ 0.05 and an absolute log2 fold change ≥ 1.

### 2.6. SOM-based transcriptional module analysis

To identify coordinated transcriptional patterns, *Vitis* genes were organized into transcriptional modules using a self-organizing map (SOM) approach. The SOM analyses was performed using oposSOM (v.2.24.0, Löffler-Wirth et al, 2015) from a gene-by-profile matrix derived from variance-stabilized (VST) expression values of the filtered *Vitis* gene-count matrix. Modules were defined from the resulting SOM landscape using D-map clustering. Each module was assigned a letter identifier and associated with its corresponding set of genes and module-level expression profile.

### 2.7. Functional enrichment of SOM modules

Gene Ontology (GO) enrichment was performed on SOM-derived transcriptional modules using topGO (v2.58.0; Alexa et al., 2022). Grapevine GO annotations were imported from the *V. vinifera* functional annotation table (https://grapedia.org) and converted into a gene-to-GO mapping. The statistical universe corresponded to the set of genes included in the SOM input matrix, while each SOM module was tested as a gene set of interest. Modules containing less than 10 genes were excluded from enrichment analysis.

For each module, GO enrichment was tested independently for Biological Process, Molecular Function and Cellular Component ontologies using Fisher’s exact tests implemented in topGO. The classic, elim, weight01 and parentchild algorithms were evaluated, with weight01 algorithm combined with Fisher’s exact test used as the primary enrichment statistic. For each module and GO, weight01 Fisher *p*-values were additionally adjusted using the Benjamini-Hochberg procedure. Terms with FDR-adjusted *p*-values ≤ 0.05 were retained enriched.

### 2.8. Transcriptome-metabolome integrative analysis

Transcriptome-metabolome integration was performed at the SOM module level to reduce transcriptomic dimensionality and relate coordinated gene expression patterns to metabolomic variation. Variance-stabilized RNA-seq expression values were obtained from DESeq2 object using the vst function. For each selected SOM module, module activity scores were calculated by first standardizing the expression of each gene across all samples (z-score) and then averaging these standardized expression values across all genes belonging to the module for each sample. Modules represented by fewer than three detected genes in the expression matrix were excluded.

Module activity scores were integrated with the curated metabolomic feature matrix described in the associated data paper. Prior to integration, transcriptomic and metabolomic variables were scaled and variables with zero variance were removed. Associations between SOM module activity and metabolite abundance were assessed using Spearman correlation coefficients. P-values were corrected for multiple testing using Benjamini-Hochberg method. Module-metabolite associations were retained when they showed an adjusted p-value <0.05 and an absolute Spearman correlation coefficient > 0.7.

## 3. RESULTS

### 3.1. Grapevine genotypes exhibit contrasted susceptibility to *Botrytis cinerea*

Contrasting levels of susceptibility to *B. cinerea* have previously been reported among grapevine genotypes, including *V. vinifera* cultivars assessed in field conditions (Dubos, 2002) or in detached-leaf infection assays (Rahman et al, 2018). Based on these data, seven *V. vinifera* cultivars (‘Cabernet Sauvignon’, ‘Chardonnay’, ‘Cot’, ‘Gamay’, ‘Merlot’, ‘Syrah’ and ‘Ugni Blanc’) were selected for a comparative susceptibility assay. Two American *Vitis* species, *V. riparia* and *V. rupestris*, commonly used as source of resistance traits in grapevine breeding (Eibach and Töpfer, 2015), were also included to broaden the range of responses to *B. cinerea* infection.

Susceptibility was first assessed using detached leaf discs inoculated with *B. cinerea* B05.10. The resulting necrotic areas revealed marked differences among grapevine genotypes (**Figure 1A**). *V. riparia* displayed the lowest symptom development under our experimental conditions, consistent with a tolerant phenotype. By contrast, extensive necrosis was observed for several *V. vinifera* cultivars, including ‘Chardonnay’, ‘Gamay’, ‘Merlot’ and ‘Cot’, as well as for *V. rupestris*, indicating higher susceptibility to *B. cinerea*. *V. vinifera* ‘Cabernet Sauvignon’, ‘Syrah’ and ‘Ugni Blanc’ showed intermediate symptom development and were therefore considered moderately susceptible in this assay.

**Figure 1.**
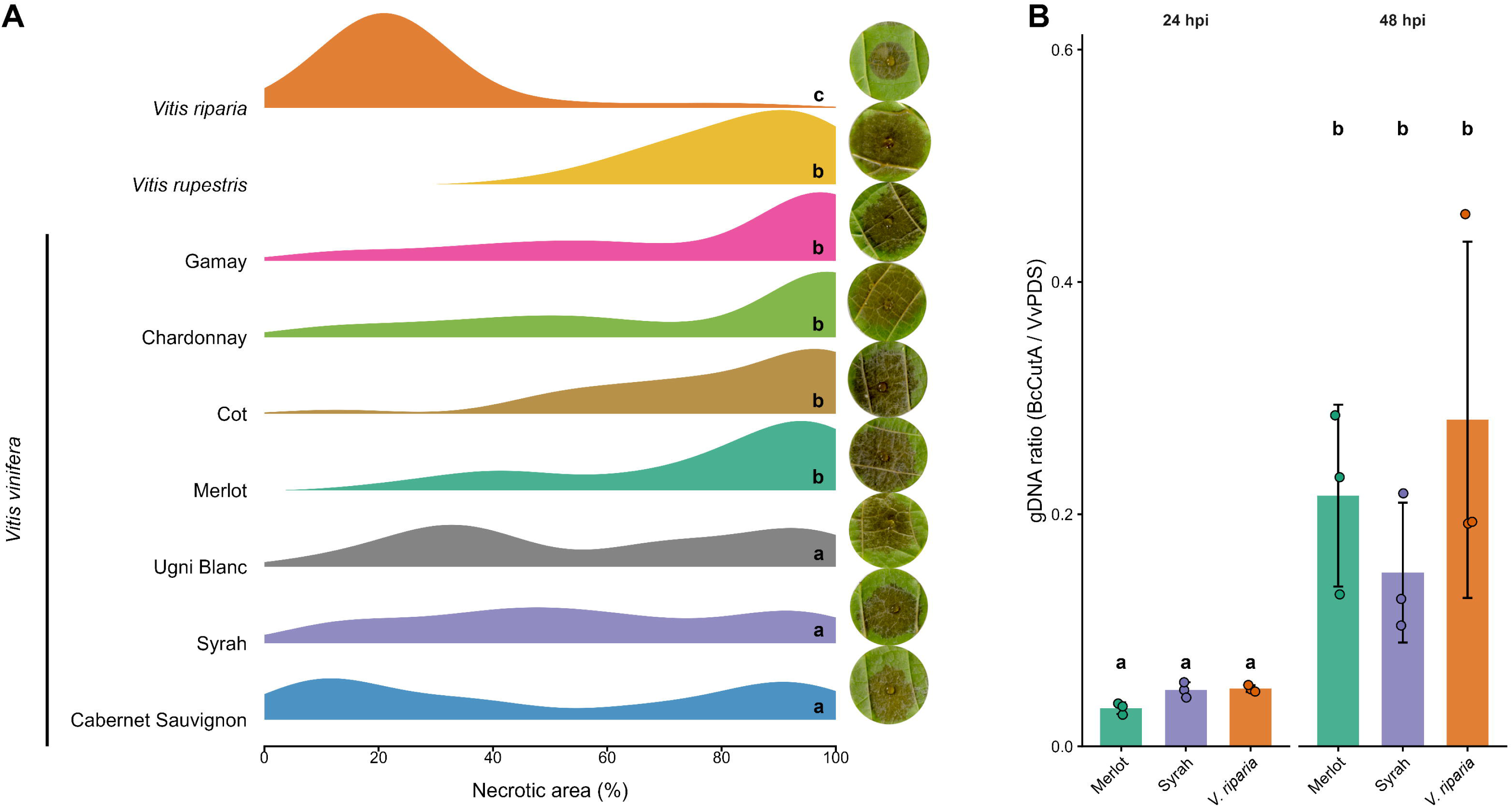
Phenotypic susceptibility and fungal biomass accumulation in grapevine genotypes infected with *Botrytis cinerea*. (A) Distribution of necrotic areas in detached leaf discs from seven <u>Vitis vinifera</u> cultivars and two American <u>Vitis</u> species inoculated with <u>B. cinerea</u> B05.10 (n=52). Necrotic areas were quantified 72 h post-inoculation using chlorophyll fluorescence imaging. Different letters indicate statistically significant differences among genotypes. Representative 2-cm-leaf-discs are shown on the left for each cultivar. (B) Quantification of <u>B. cinerea</u> biomass in infected leaves of <u>V. vinifera</u> ‘Merlot’ and ‘Syrah’, and <u>V. riparia</u> at 24 and 48 h post-infection (n=3). Fungal biomass was estimated by qPCR as the ratio of <u>B. cinerea</u> to grapevine genomic DNA using BcCutA and VvPDS genes respectively. Different letters indicate a significant overall effect of sampling time across genotypes, as determined using an aligned rank transform model (p-value < 0.05).

This phenotypic screening confirmed a broad variation in susceptibility to *B. cinerea* among *Vitis* genotypes and supported the selection of three representative genotypes for omic analyses: *V. riparia* as the genotype displaying the lowest symptom development, *V. vinifera* ‘Merlot’ as a highly susceptible cultivar, and *V. vinifera* ‘Syrah’ as a moderately susceptible cultivar.

To determine whether the ranking of symptom severity was paralleled by differences in fungal DNA accumulation, *B. cinerea* biomass was quantified in infected leaves of the three selected genotypes at 24 and 48 hpi using qPCR (**Figure 1B**). The ratio of fungal to grapevine genomic DNA increased between 24 and 48 hpi in all three genotypes, confirming fungal progression during infection. The aligned rank transformed model indicated a non-significant genotype effect (F = 1.0125; df = 2; p = 0.39), a significant time effect (F = 38.1453; df = 1; p = 0,000048) and a non-significant time x genotype interaction (F = 1.5133; df = 2; p = 0.26). Fungal DNA accumulation did not mirror the symptom-severity ranking. These results indicate that the contrasted symptom phenotypes observed among the three genotypes cannot be explained solely by differences in fungal growth. We therefore used transcriptomic and metabolomic analyses to investigate the host molecular responses underlying these contrasted susceptibility levels.

### 3.2. *B. cinerea* infection triggers extensive and time-dependent transcriptional reprogramming in grapevine

To characterize transcriptional responses of grapevines to *B. cinerea*, differential expression analyses were performed independently for each genotype by comparing infected leaves with their corresponding mock-treated controls at 24 and 48 hpi (**Supplementary Table S2**). Infection induced a large-scale reprogramming of the *Vitis* transcriptome in all three genotypes, with thousands of genes differentially expressed at both time points (**Figure 2**). At 24 hpi, the number of differentially expressed genes ranged from 4 306 in *V. riparia* to 6 118 in *V. vinifera ‘*Syrah’, with *V. vinifera* ‘Merlot’ showing an intermediate number of 5 337 DEGs (**Figure 2A**). The response was further amplified at 48 hpi, reaching 13 376 DEGs in *V. riparia*, 13 758 in *V. vinifera* ‘Merlot’ and 14 399 in *V. vinifera* ‘Syrah’ (**Figure 2B**).

**Figure 2.**
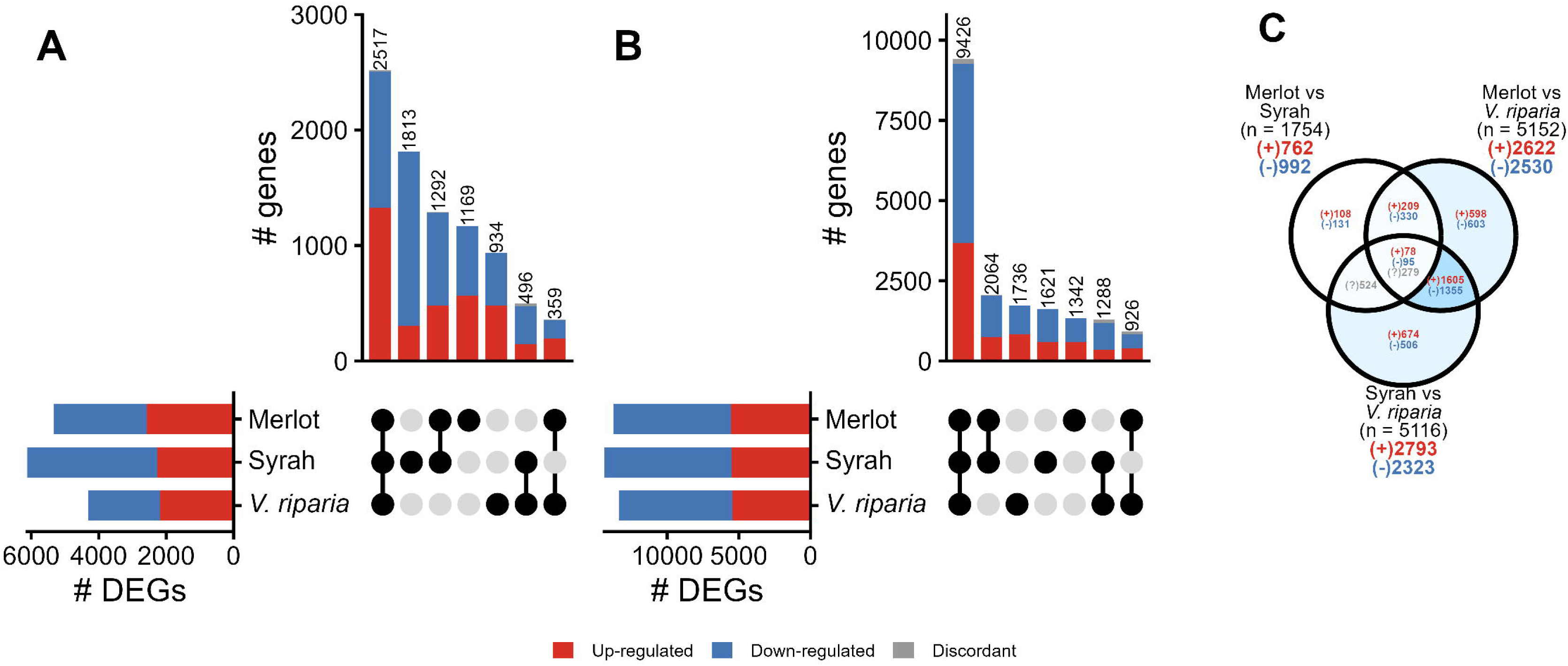
Overlap of differentially expressed genes (DEGs) among grapevine genotypes during *Botrytis cinerea* infection and under non-treated conditions (A–B) UpSet plots showing the overlap of DEGs identified in <u>V. vinifera</u> ‘Merlot’ and ‘Syrah’ and <u>V. riparia</u> at 24 hpi (A) and 48 hpi (B) following <u>B. cinerea</u> infection, relative to the corresponding mock-treated samples. Horizontal bars indicate the total number of DEGs detected in each genotype and their distribution between up- and down-regulated genes. Vertical bars indicate the number of genes belonging to each genotype-specific or shared interactions. Genes consistently regulated in the same direction across genotypes represented in an intersection are classified as up- or down-regulated, whereas genes displaying opposite directions of regulation among genotypes are classified as discordant. Only non-empty intersections are shown. (C) Venn diagram showing the overlap of DEGs obtained from the three pairwise comparisons between non-treated <u>V. vinifera</u> ‘Merlot’ and ‘Syrah’, and <u>V. riparia</u> samples. The total number of DEGs detected in each pairwise comparison is indicated under brackets. (+) indicates genes more highly expressed in the first genotype of the comparison, (-) indicates genes more highly expressed in the second genotype, and (?) indicates discordant direction among shared comparisons. In all panels, only grapevine genes were considered. DEGs were defined using an adjusted P-value < 0.05 and |log2 fold change| > 1.

The distribution of DEGs among genotypes revealed both shared and genotype-associated components of the infection response. At 24 hpi, 2 517 DEGs were common to all three genotypes, corresponding to 29.3% of the infection-responsive genes at this time point (**Figure 2A**). This common set represented 47.2% of Merlot DEGs, 41.1% of Syrah DEGs and 58.5% of *V. riparia* DEGs. At 48 hpi, the shared response became more prominent, with 9 426 DEGs common to the three genotypes, representing 51.2% of the infection-responsive genes at this time point and 65.5-70.5% of the DEGs detected within each genotype (**Figure 2B**). Thus, although genotype-specific transcriptional responses were observed, particularly at 24 hpi, the late response to *B. cinerea* was largely dominated by a conserved transcriptional reprogramming shared by the three genotypes.

Directionality analysis further indicated that this common response was highly consistent among genotypes (**Figure 2A-B**). Among the DEGs shared by the three genotypes, only 9 genes at 24 hpi and 152 genes at 48 hpi showed discordant regulation directions. Most shared DEGs were therefore in the same direction across the three genotypes. At 24 hpi, the common response included both induced and repressed genes, whereas at 48 hpi the shared response was dominated by a large set of commonly repressed genes together with a substantial set of commonly induced genes.

Basal transcriptional differences were also detected before infection (**Figure 2C**). In non-treated leaves, 5 341 genes were differentially expressed between *V. riparia* and at least one *V. vinifera*, while not being differentially expressed between *V. vinifera* ‘Merlot’ and ‘Syrah’. These genes included 2 877 genes expressed at higher levels in *V. riparia* and 2 464 genes expressed at lower levels. These basal differences indicate that *V. riparia* was already transcriptionally distinct from *V. vinifera* before pathogen challenge.

### 3.3. Sample-level SOM portraits reveal reproducible transcriptional states and genotype-specific infection trajectories

To further explore the structure of the transcriptomic dataset beyond individual differential expression contrasts, a SOM analysis was applied to the filtered *Vitis* expression matrix. SOM-based transcriptome portrayal has recently proved useful in grapevine for resolving sample-specific expression landscapes and identifying coordinated stress-responsive gene modules across cultivars and environmental conditions (Konecny et al, 2023). This approach projected each sample onto an expression portrait, allowing global transcriptional states to be compared across genotypes, treatments and time points. The sample-level SOM portraits showed highly reproducible expression patterns among biological replicates within each genotype-by-condition combination (**Supplementary Figure S1**). This reproducibility was observed across non-treated, mock-treated and infected samples, supporting the consistency of the RNA-seq dataset and the robustness of the experimental design.

The SOM portraits also revealed genotype- and time-dependent infection pattern (**Figure 3**). In non-treated conditions, *V. riparia* displayed a transcriptional landscape distinct from those of *V. vinifera* ‘Merlot’ and ‘Syrah’, consistent with the basal differences detected by differential expression analysis. At 24 hpi, infected *V. riparia* samples showed a specific expression portrait that differed from both its mock-treated controls and the infected *V. vinifera* samples. By contrast, at 48 hpi, infected samples from the three genotypes displayed more similar global SOM patterns, suggesting convergence toward a shared late infection-associated transcriptional state. This convergence was accompanied by low sample-specific silhouette values, indicating that late infected samples were less clearly separated according to genotype than earlier or non-infected states.

**Figure 3.**
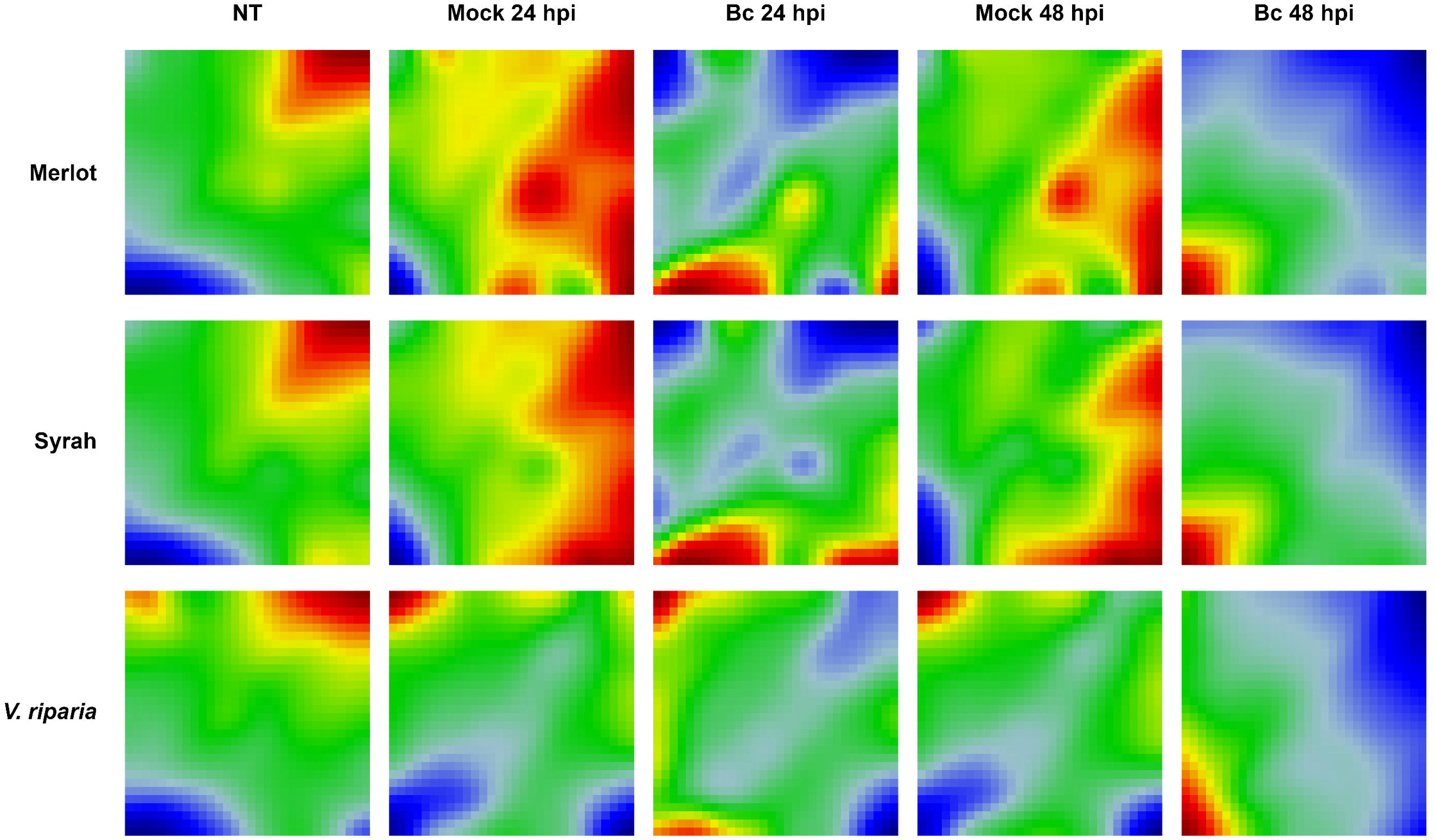
Average self-organizing map (SOM) expression portraits across grapevine genotypes and experimental conditions Average SOM portraits are shown for <u>V. vinifera</u> ‘Merlot’ and ‘Syrah’ and <u>V. riparia</u> under non-treated conditions (NT), mock treatment at 24 and 48 hpi, and <u>B. cinerea</u> (Bc) infection at 24 and 48 hpi. Each portrait represents the mean metagene expression profile calculated from the three biological replicates for a given genotype x condition combination. Colors indicate relative metagene expression within each portrait, from low expression (blue) to high expression (red). Rows correspond to grapevine genotypes and columns to experimental conditions.

### 3.4. Contrast-based SOM clustering resolves basal, shared and genotype-associated transcriptional trajectories

To summarize the major transcriptional trajectories underlying the response to *B. cinerea*, a contrast-based SOM analysis was performed on the subset of genes contributing to the main expression changes across the experiment. D-map clustering of the contrast landscape identified 14 transcriptional modules (A to N), comprising 8 990 genes in total (**Figure 4A; Supplementary Table S2**). Module size varied markedly, ranging from 109 genes in module A to 2 620 genes in module M (**Figure 4B**). Six modules (B, C, H, J, L, and N) displayed profiles of particular interest with respect to basal genotype differences or responses to *B. cinerea* infection. The contribution of infection-responsive genes to these modules was further assessed by intersecting module membership with the differential expression analysis (**Figure 4B**).

**Figure 4.**
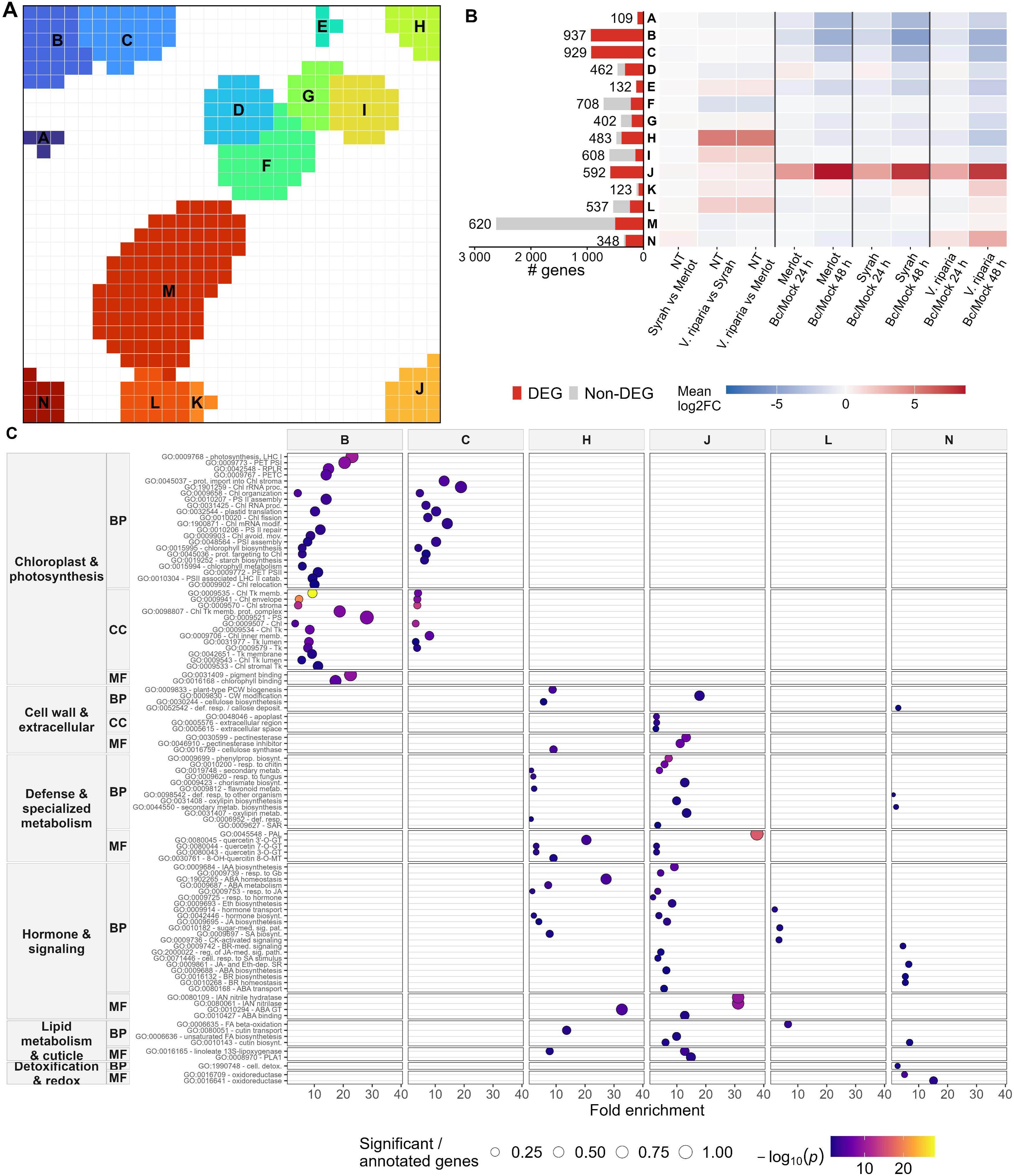
D-map clustering, differential expression profiles and functional enrichment of SOM-derived transcriptional modules (A) D-map representation of the contrast-based self-organizing map, showing the 14 transcriptional modules A to N identified from the 30 x 30 metagene grid. Colors distinguish individual D-clusters and do not represent expression levels. (B) Composition and differential expression profiles of the 14 SOM module. Left, number of genes assigned to each module. Red bar indicates genes identified as differentially expressed in at least one <u>B. cinerea</u> versus mock comparison at 24 or 48 hpi, whereas gray bars represent non-DEGs. Numbers indicate the total number of genes in each module. Right, differential expression profile across genotypes and infection contrasts. For non-treated samples (NT), pairwise comparisons correspond to Syrah versus Merlot, <u>V. riparia</u> vs Merlot and <u>V. riparia</u> vs Syrah. Infection-associated responses correspond to <u>B.</u> <u>cinerea</u>-infected vs time-matched mock-treated samples at 24 and 48 hpi for each genotype. Each cell represents the mean DESeq2 log2 fold change of genes assigned to the corresponding module. Red indicates positive and blue negative differential expression, with color intensity proportional to the magnitude of the mean log2 fold change. Vertical lines separate the NT comparisons from the Merlot, Syrah and <u>V. riparia</u> infection-response contrasts. (C) Functional enrichment of selected SOM modules. Enriched gene ontology (GO) terms were manually organized into six functional groups: chloroplast and photosynthesis, cell wall and extracellular processes, defense and specialized metabolism, hormone and signaling, lipid metabolism and cuticle, and detoxification and redox processes. Within each functional group, GO terms are further separated according to GO ontology: biological process (BP), cellular component (CC) and molecular function (MF). Columns correspond to SOM modules, while rows represent individual GO terms, displayed as GO identifier followed by a shortened term description. Dot position indicates fold enrichment, dot size represents the proportion of significant genes among genes annotated to the corresponding GO term, and dot color represents enrichment significance as −log10(p−value). Only significantly enriched terms selected for their relevance to the biological signatures of the respective modules are shown.

Module B (937 genes) and C (929 genes) showed a similar pattern, characterized by a marked decrease in expression at 48 hpi in all three genotypes (**Figure 4B**). These modules contained among the highest proportion of infection-responsive genes. At 48 hpi, all 937 genes of module B were down-regulated in both *V. vinifera* cultivars. In *V. riparia*, 929 genes were differentially expressed, including 928 down-regulated genes. Module C showed a similarly homogeneous response: 920 of 929 genes were down-regulated in *V. vinifera* ‘Merlot’, while all 929 genes were down-regulated in *V. vinifera* ‘Syrah’ and *V. riparia*. Conversely, module J (592 genes) showed a strong induction following infection. At 24 hpi, 83.6-88.3% of its genes were differentially expressed across genotypes, predominantly showing increased expression. At 48 hpi and 99.7-100% of module J genes were differentially expressed, all being up-regulated. Several functionally characterized grapevine genes discussed below followed this module-J trajectory, including *VvMYB14* (Vitvi05_01chr07g09050), *VvSTS36* (Vitvi05_01chr16g14430), *VvSTS39* (Vitvi05_01chr16g14470), *VvSTS41* (Vitvi05_01chr16g14490), *VvABCB15* (Vitvi05_01chr14g28020), *VvHT5* (Vitvi05_01chr05g07090) and *VvPME10* (Vitvi05_01chr06g16750) (**Supplementary Figure S2**). Notably, the module-J *VvMYB14, VvSTS36* and *VvSTS39* also showed higher basal expression in *V. riparia* than at least one of the two *V. vinifera* cultivars, indicating that this module combines basal genotype-associated differences with a strong infection-responsive component (**Supplementary Table S2**).

Module H (483 genes) and L (537 genes) primarily distinguished *V. riparia* from the two *V. vinifera* cultivars under non-treated conditions. In module H, 449 of the 483 genes were differentially expressed between *V. riparia* and *V. vinifera* ‘Merlot’ and 448 between *V. riparia* and *V. vinifera* ‘Syrah’ under non-treated conditions. All of these DEGs showed higher expression in *V. riparia*. In addition to this basal pattern, module H expression decreased markedly at 48 hpi, with 68.7% of its genes differentially expressed in *V. riparia*, compared with 35.6% in Merlot and 43.5% in Syrah, most of them being down-regulated. Module L displayed a similar but less pronounced pattern, with 198 of 537 genes differentially expressed between *V. riparia* and *V. vinifera* ‘Merlot’, and 171 between *V. riparia* and *V. vinifera* ‘Syrah’. Of these, 198 and 169 genes, respectively, showed higher expression in *V. riparia*.

Finally, module N (348 genes) showed a distinct genotype-dependent infection response that was already detectable at 24 hpi and became more pronounced at 48 hpi. At 24 hpi, 19.3%, 21% and 26.1% of module N genes were differentially expressed in *V. vinifera* ‘Merlot’, ‘Syrah’ and *V. riparia*, respectively. At 48 hpi, the proportion increased to 43.7% in *V. vinifera* ‘Merlot’, 44.3% in *V. vinifera* ‘Syrah’ and 67.2% in *V. riparia*. In *V. riparia*, 233 of the 234 DEGs were up-regulated, whereas DEGs in both *V. vinifera* cultivars were predominantly down-regulated (132/152 and 124/154, respectively). Among the genes belonging to module N, a putative respiratory burst oxidase homolog (RBOH, Vitvi05_01chr02g00560) was significantly induced in *V. riparia* at both 24 and 48 hpi, whereas it was not identified as differentially expressed in either *V. vinifera* cultivars under the applied DEG criteria (**Supplementary Figure S2**).

### 3.5. Functional enrichment of selected SOM modules highlights plastidial, cell wall, hormone and defense-related processes

Functional enrichment analysis revealed distinct biological signatures among the selected SOM modules (**Figure 4C, Supplementary Table S3**). Modules B and C were predominantly associated with chloroplast functions. Module B was strongly enriched in photosynthesis-related processes, including light harvesting, photosynthetic electron transport, photosystem II repair and chloroplast organization. Consistently, chlorophyll and pigment-binding activities and chloroplast thylakoid, envelope and stroma compartment function were overrepresented. Module C was characterized by chloroplast organization, protein import into the chloroplast, chloroplast RNA processing, plastid translation and photosystem I assembly, together with strong enrichment of chloroplast stroma, envelop and thylakoid compartment functions. Starch biosynthesis was also represented.

Module H was enriched in primary cell wall biogenesis and cellulose biosynthesis, including cellulose synthase activity. Hormone-related terms included abscisic acid homeostasis and metabolism and response to jasmonic acid. Defense response and response to fungus were also enriched, together with quercetin glucosyltransferase activities and cutin transport. Module L was mainly associated with fatty acid β-oxidation, with additional enrichment in hormone transport, sugar-mediated signaling and cytokinin signaling.

Module J displayed a broad defense-associated functional signature. Phenylpropanoid biosynthesis and phenylalanine ammonia-lyase activity were among the strongest enrichments, together with chitin and specialized metabolism. Consistent with this signature, module J contained several characterized components of grapevine stilbenoid metabolism, including the transcriptional regulator *VvMYB14* and the stilbene synthase genes *VvSTS36*, *VvSTS39* and *VvSTS41* **(Supplementary Figure S2)**. The functionally related regulator *VvMY15* (Vitvi05_01chr05g11030) and the class III peroxidase *VvPRX4* (Vitvi05_01chr06g11220), although not assigned to a D-map module, also showed infection-responsive expression patterns **(Supplementary Figure S2)**. Module J additionally contained *VvABCB15*, encoding a transporter subsequently discussed in relation to resveratrol export, which showed a marked induction at 48 hpi in all three genotypes **(Supplementary Figure S2)**. Beyond these specialized-metabolism components, the functional signature of module J extended to multiple signaling, lipid-metabolism and cell-wall-related processes. Hormone-related terms included responses to jasmonic acid and gibberellin, ethylene biosynthesis and abscisic acid-related processes. Lipid-related function included linoleate 13S-lipoxygenase and phospholipase A1 activities, unsaturated fatty acid and cutin biosynthesis. Pectinesterase and pectinesterase inhibitor activities were also strongly enriched, together with apoplastic and extracellular localization. This cell -wall-related signature included *VvPME10*, which belongs to module J and was strongly induced following infection in all three genotypes **(Supplementary Figure S2)**. Its functionally characterized transcriptional regulator *VvWRKY03* (Vitvi05_01chr01g23290), although not assigned to a D-map module, was likewise induced at both 24 and 48 hpi in the three genotypes **(Supplementary Figure S2)**.

Module N was enriched in defense response to other organisms and secondary metabolite biosynthesis. Additional terms included brassinosteroid signaling, biosynthesis and homeostasis, cellular detoxification, cutin biosynthesis, jasmonic acid- and ethylene-dependent systemic resistance, and defense response by callose deposition.

### 3.6. SOM-metabolite correlations associate transcriptional module with distinct metabolic changes

To extend the transcriptional analysis to the metabolic dimension of the response to *B. cinerea*, untargeted metabolomic profiling was performed on the same biological samples used for RNA-seq. Metabolomic profiles were primarily structured by genotype, with Merlot, Syrah and *Vitis riparia* displaying distinct metabolic signatures (Monnereau et al, *submitted to Data in Brief*). At 48 hpi, infected samples underwent a marked metabolic shift, while retaining a clear genotype-dependent structure. This paired design enabled transcriptional programs identified by SOM analysis to be associated with metabolite accumulation patterns across genotypes, treatment and time points. We therefore integrated module-level expression profiles with metabolite abundance to determine whether the major basal and infection-responsive trajectories were accompanied by metabolic changes. Correlation analysis identified significant associations between metabolite abundance and module B, C, H and J, whereas no association reached the selected correlation threshold for modules L and N (**Figure 5A, Supplementary Table S4**).

**Figure 5.**
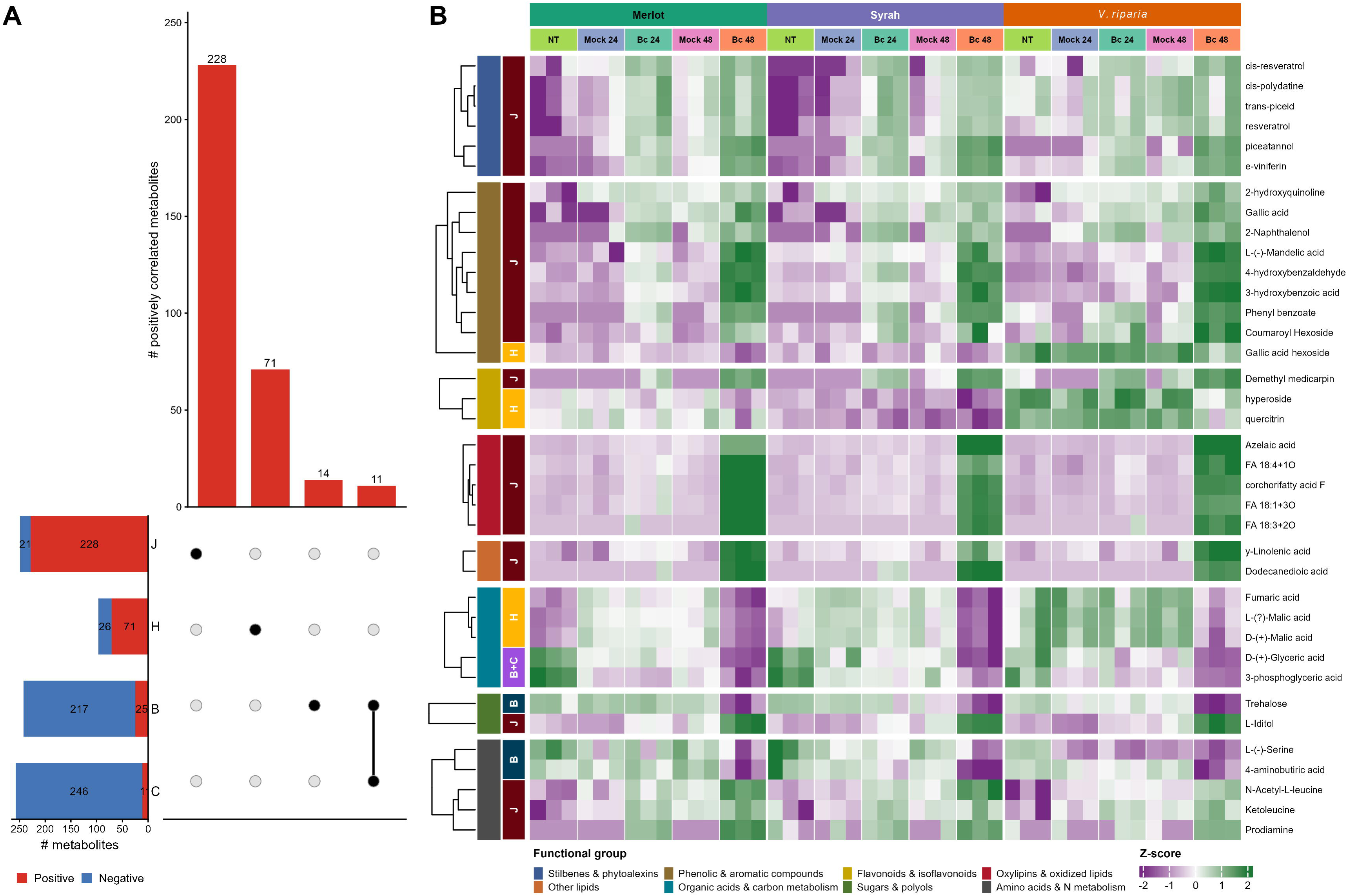
Association between SOM transcriptional modules and metabolite accumulation profiles (A) upset representation of significant correlations between metabolite abundance and SOM modules B, C, H and J. Horizontal set-size bar shows the total number of significantly correlated metabolites for each module, separated into positive (red) and negative (blue) correlations. Vertical bars and the intersection matrix represent only positively correlated metabolites and indicate the number of metabolites uniquely or jointly associated with different SOM modules. (B) Heatmap showing the accumulation profiles of MS/MS-supported metabolites positively correlated with at least one of the selected SOM modules. When several features corresponded to the same metabolite and displayed the same module-association pattern, their standardized abundance profiles were averaged and represented as a single row. Metabolites are grouped according to functional classes, indicated by the first row annotation, while the second row annotation indicates the SOM module(s) with which each metabolite is positively correlated. Columns correspond to biological replicates from <u>V. vinifera</u> ‘Merlot’ and ‘Syrah’ and <u>V. riparia</u> under non-treated (NT), mock-treated and <u>B. cinerea</u>-infected conditions (Bc) at 24 and 48 hpi. Colors in the heatmap represent row-wise standardized metabolite abundance (Z-score), from lower (blue) to higher (red) relative accumulation. Only metabolites assigned to the displayed functional groups and supported by MS/MS annotation are shown.

Module B and C showed predominantly negative metabolite correlations (217/242 and 246/257, respectively). Conversely, several metabolites related to primary carbon metabolism were positively correlated with these modules. In module B, these included 3-phosphoglyceric acid (3-PGA), glyceric acid, serine, Gamma-aminobutyric acid (GABA) and trehalose, together with sucrose and galactinol among features without MS/MS confirmation. These metabolites followed a comparable profile, with higher accumulation in non-infected samples and reduced abundance during infection, particularly at 48 hpi (**Figure 5B)**. Module C was similarly positively associated with 3-PGA and glyceric acid.

Module H showed 71 positive and 26 negative metabolite associations. Positive correlations included several malic acid features, fumaric acid, gallic acid hexoside and the flavonols hyperoside and quercitrin, defining a metabolic profile dominated by organic acids and glycosylated phenolic compounds. These metabolites were generally more abundant in *V. riparia* under non-treated conditions but showed very low accumulation at 48 hpi in all three genotypes, with the largest decrease observed in *V. riparia* (**Figure 5B)**.

Module J showed predominantly positive associations, with 228 of 249 correlated features displaying positive correlations. These included several stilbenes, such as trans-ε-viniferin, resveratrol, cis-resveratrol, piceatannol and trans-piceid, together with gallic acid, 4-hydroxybenzaldehyde, coumaroyl hexoside and demetylmedicarpin. Positive correlations were also detected for oxidized C18 fatty acids, γ-linolenic acid and azelaic acid. Features annotated without MS/MS further included α-linolenic acid, jasmonic acid and jasmonoyl-isoleucine. Most of these metabolites showed low basal abundance and accumulated following infection, with the strongest increase observed at 48 hpi in all three genotypes (**Figure 5B)**.

Together, these associations show that the major SOM-derived transcriptional programs were accompanied by coherent metabolic changes, linking basal and infection responsive gene-expression trajectories to distinct shifts in primary carbon metabolism, phenolic compounds and lipid-derived metabolites.

## 4. DISCUSSION

### 4.1. A preconfigured basal state and an early divergent response underpin reduced symptoms in *Vitis riparia*

The reduced lesion development of *V. riparia* was not accompanied by lower fungal DNA accumulation at 48 hpi, suggesting that its phenotype cannot be explained solely by enhanced restriction of *B. cinerea* growth. This distinction is important because resistance and tolerance represent different components of plant defense: resistance limits pathogen multiplication, whereas tolerance reduces the damage caused by infection at a given pathogen burden (Pagán and García-Arenal, 2020). Although formally demonstrating tolerance requires quantifying the relationship between pathogen burden and host damage, the different rankings obtained for fungal DNA and symptom development are compatible with a disease tolerance component in *V. riparia*.

This phenotype was associated with molecular differences that preceded the large transcriptional response observed at 48 hpi. *V. riparia* was already distinct from both *V. vinifera* cultivars before infection and displayed a specific transcriptional state at 24 hpi. A similar importance of pre-infection properties and early responses was reported in the resistant wild grapevine *V. amurensis* in which resistance to *B. cinerea* was associated with basal differences in antioxidant, carbon/nitrogen and defense metabolism together with earlier cell-wall responses (Wan et al, 2021). Interestingly, this differs from the *V. riparia*-*Plasmopara viticola* interaction, where resistance was predominantly associated with rapid post-infection reprogramming rather than constitutively higher expression of defense genes (Polesani et al, 2010), suggesting that the contribution of basal preconfiguration is pathogen-dependent. The distinctive response observed here at 24 hpi therefore points to the early, largely asymptomatic phase as a potentially critical period determining subsequent symptom development. Such a timing is compatible with the transient asymptomatic or biotrophic-like phase proposed during the establishment of *B. cinerea* infection before extensive necrotrophic growth, although the trophic nature of this early phase remains debated (Rahman et al, 2024).

Modules H and L provide potential molecular components of this preconfigured state, including differences in cell-wall biogenesis, cellulose synthesis, lipid metabolism and hormone-related processes. These transcriptional differences were accompanied by higher basal accumulation in *V. riparia* of organic acids and glycosylated phenolics, including gallic acid hexoside, hyperoside and quercitrin. Constitutively accumulated specialized metabolites with antimicrobial or protective functions are commonly referred to as phytoanticipins, in contrast to phytoalexins synthesized or strongly induced after pathogen perception (Piasecka et al, 2015; Pedras and Yaya, 2015). Glycosylation is particularly relevant to such performed chemical defenses because it increases solubility and stability and allows potentially bioactive compounds to be stored in less reactive forms that can subsequently be mobilized following stress (Le Roy et al, 2016). Accordingly, the higher basal pool of glycosylated phenolics in *V. riparia* may represent a phytoanticipin-like chemical reservoir rather than simply a constitutive elevation of inducible defense metabolism.

The marked depletion of these compounds at 48 hpi suggests substantial turnover of this basal pool during infection. Deglycosylation and subsequent metabolism of stored phenolics provide one possible mechanism, as reversible glycosylation is widely used by plants to regulate the availability of specialized metabolites during defense (Morant et al, 2008; Zhang et al, 2022). Their decline could also result from oxidation. *B. cinerea* secretes laccases capable of oxidizing a broad range of grape phenolics, and infection-dependent losses of phenolic compounds have been linked to laccase activity in grape tissues (Zimdars et al, 2017; Umberath et al, 2024, 2026). These compounds should therefore not be considered direct precursors of the stilbenoids accumulating later during infection. Rather, their depletion, concomitant with the activation of inducible phenylpropanoid metabolism, supports a broader transition from a pre-existing phenolic pool towards an infection-induced chemical defense state. This transition may also be facilitated by the basal transcriptional state. *VvMYB14* and *VvMYB15*, experimentally characterized activators of grapevine stilbene synthase (*STS*) genes, were more highly expressed in untreated *V. riparia*, as were *VvSTS36* and *VvSTS39* (Höll et al, 2013). Thus, the distinct pre-infection state could combine an existing pool of glycosylated phenolics with greater transcriptional preparedness for subsequent stilbenoid production. The latter interpretation concerns regulatory potential, not demonstrated constitutive accumulation of stilbenes.

### 4.2. A conserved late response to *B. cinerea* involves a profound reorganization of chloroplast and photosynthetic functions

The predominance of shared DEGs at 48 hpi indicated that the late response to *B. cinerea* is largely conserved across the three grapevine genotypes. A major component of this common response was the coordinated repression of module B and C, which encompass photosynthesis, photosystem organization, chloroplast biogenesis and plastid-associated functions. The concomitant decrease in 3-PGA, glycerate and other carbon-related metabolites positively associated with these modules further indicates that transcriptional repression was accompanied by substantial metabolic reorganization. A similar repression of photosynthesis-related transcriptional programs has repeatedly been reported during *B. cinerea* infection. In *Arabidopsis thaliana*, high-resolution transcriptomics revealed coordinated repression of chlorophyll biosynthesis, photosystems I and II and chloroplast organization from the early stages of infection onwards (Windram et al 2012). Photosynthesis-related genes are also repressed in *B. cinerea*-infected grape berries (Agudelo-Romero et al, 2015) and leaves of both *V. vinifera* and *V. amurensis* (Wan et al, 2021). More broadly, repression of photosynthetic gene expression is a recurrent feature of plant responses to diverse biotic stresses (Bilgin et al, 2010; Cheaib and Killiny, 2025).

This response has often been interpreted within the framework of a growth-immunity trade-off, whereby resources normally invested in photosynthetic activity and growth are redirected towards the synthesis of defense proteins and specialized metabolites (Bilgin et al, 2010). However, chloroplast reorganization during infection likely extends beyond passive resource reallocation. Chloroplasts are major immune-signaling hubs involved in reactive oxygen species (ROS) and Ca^2+^ signaling, chloroplast-to-nucleus retrograde communication and the biosynthesis of defense-related hormones, including jasmonic acid, salicylic acid and abscisic acid (Rui et al, 2025; Sun et al, 2025). They also contribute precursors for cuticular and cell-wall-associated defenses (Sun et al, 2025). The relationship between chloroplast activity and immunity is particularly complex during infection by necrotrophs. In tobacco, reducing chloroplast-derived ROS decreased both tissue damage and fungal growth during *B. cinerea* infection, demonstrating that chloroplastic ROS can contribute to disease development rather than exclusively host defense (Rossi et al, 2017). Thus, the strong repression of modules B and C is more consistent with a broad functional reconfiguration of chloroplast activity than with a repression of photosynthesis-associated gene expression.

Such reconfiguration may also connect the decline in photosynthetic carbon metabolism with the defense pathways activated during late infection. Chloroplasts are the primary site of fatty-acid synthesis and provide lipid substrates for oxylipin and jasmonate production, while also contributing to the synthesis of surface barriers such as cutin and waxes (Sun et al, 2025). The coordinated repression of photosynthetic/plastidial functions and activation of lipid-, hormone-, and defense-related processes observed in module J may therefore reflect a broad redistribution of metabolic function during infection, although carbon flux cannot be inferred directly from transcript abundance. Not all components of this common response are necessarily protective. *VvHT5*, a hexose transporter strongly induced at 48hpi in all three genotypes, was recently shown to increase susceptibility to *B. cinerea* when overexpressed in grapevine (Monnereau et al, 2026). This shared late response may encompass both defense-associated metabolism and processes exploitable by the pathogen. It also helps explain why similar transcriptional responses can accompany different disease outcome.

### 4.3. Phenylpropanoid and stilbenoid metabolism constitutes a major conserved defense response to *B. cinerea*

Module J identifies phenylpropanoid metabolism as one of the major conserved responses to *B. cinerea* infection. Its strong enrichment in phenylalanine ammonia-lyase activity and phenylpropanoid biosynthesis was accompanied by the accumulation of several stilbenoids, including resveratrol, piceatannol, piceid and ε-viniferin derivatives. This coordinated transcript-metabolite response is supported at gene level by the induction of *VvSTS41* alongside its experimentally characterized transcriptional regulators *VvMYB14* and *VvMYB15* (Höll et al, 2013). It is also consistent with previous grapevine studies showing that *B. cinerea* induces PAL- and stilbene synthase-associated pathways together with increased accumulation of trans-resveratrol and other phenolic defense metabolites (Agudelo-Romero et al, 2015; Maia et al., 2023). Developmental studies of the grapevine-*B. cinerea* interaction have likewise shown that stilbene-associated defenses vary strongly with berry developmental stage and susceptibility (Kelloniemi et al, 2015), while the broader regulation and diversification of stilbene biosynthesis across grapevine species have been extensively documented (Dubrovina and Kiselev, 2017). In infected *V. vinifera* ‘Trincadeira’ berries, for instance, activation of phenylalanine and phenylpropanoid metabolism coincided with the induction of several PAL and stilbene synthase genes and increased trans-resveratrol accumulation (Agudelo-Romero et al, 2015). Stilbenoids are among the best-characterized inducible phytoalexins of grapevines. Their defensive relevance is supported both by their accumulation at infection sites and by direct antifungal activity against grapevine pathogens. In particular, resveratrol and oligomerized stilbenes such as ε-viniferin inhibit *B. cinerea* mycelial growth and conidial germination, with complex stilbene mixtures retaining marked antifungal activity in vitro (Adrian et al, 1997; Taillis et al., 2023). Beyond stilbene biosynthesis, our transcriptomic data also point to the activation of enzymatic processes involved in their structural diversification. The gene encoding VvPRX4, a class III peroxidase recently shown to catalyze resveratrol oligomerization and the conversion of ε-viniferin into higher-order oligomers *in vitro* (Marant et al, 2025), was significantly induced at both 24 and 48 hpi in all three genotypes. Together with the accumulation of resveratrol and ε-viniferin-related compounds, this response is consistent with the involvement of host peroxidases in infection-associated stilbene metabolism, although the contribution of *VvPRX4* to the formation of individual oligomers *in planta* remains to be established. Such oligomerization has long been associated with grapevine chemical defense, with viniferins generally displaying stronger antifungal activity than resveratrol itself (Langcake, 1981; Pezet et al., 2003a; Taillis et al., 2023).

However, the strong activation of module J in all three genotypes indicates that induction of the stilbenoid pathway *per se* does not distinguish the reduced-necrosis phenotype of *V. riparia* from the two *V. vinifera* at this late stage. This does not imply equivalent defensive effectiveness, which may depend on the abundance, localization, structural diversification and biological activity of the resulting stilbenoids. This distinction is consistent with earlier studies showing that induction of resveratrol does not necessarily predict *B. cinerea* resistance. In a broad survey of grape genotypes, resveratrol accumulation after inoculation was detected mainly in susceptible or moderately resistant accessions and was not correlated with resistance *per se* (Gabler et al., 2003). Likewise, the susceptible cultivar *V. vinifera* ‘Trincadeira’ mounts a pronounced phenylpropanoid/stilbenoid response after *B. cinerea* infection without preventing disease progression (Agudelo-Romero et al., 2015). Thus, stilbenoid induction appears to constitute a core grapevine response to *B. cinerea*, whereas differences in disease outcome are more likely to depend on timing, metabolic context of additional defense programs in which this response is embedded.

The effectiveness of this pathway may also depend on where the accumulated compounds are delivered and how they are transformed. The strong induction of *VvABCB15* in module J is relevant in this regard because this transporter has been shown to export trans-resveratrol from grapevine cells (Martinez-Marquez et al, 2025). Although its transport activity was not assessed here, this points to spatial availability as a potential determinant of phytoalexin action. Moreover, *B. cinerea* can metabolize stilbenes through its secretome: resveratrol and ε-viniferin can be converted into more complex dimers and tetramers, while other oligomers are more resistant to fungal transformation (Gindro et al, 2017; Taillis et al., 2023). Earlier work also showed that *B. cinerea* can oxidize resveratrol through laccase-like stilbene-oxidase activity (Adrian et al, 1998). These observations reinforce the idea that disease outcome cannot be inferred from pathway activation alone, but likely depends on the timing of phytoalexin production, their localization, structural diversification and subsequent biotransformation by the fungus (Gindro et al, 2017; Taillis et al, 2023).

### 4.4. Lipid reprogramming may connect defense signaling with reinforcement of surface barriers

The lipid-related signature of module J suggests that lipid metabolism participates in the conserved response to *B. cinerea* at both signaling and structural levels. Enrichment in linoleate 13S-lipoxygenase and phospholipase A1 activities, together with unsaturated fatty-acid biosynthesis, was accompanied by positive association with oxidized C18 fatty acids and γ-linolenic acid. The additional detection of α-linolenic acid, jasmonic acid and jasmonoyl-isoleucine among features lacking MS/MS confirmation further supports activation of an oxylipin-related pathway. This is consistent with the established role of C18 fatty acid as substrates for lipoxygenase-dependent oxylipin and jasmonate synthesis and with the importance of jasmonic acid signaling in defense against necrotrophic pathogens (La Camera et al, 2004). Consistent with our observations, *B. cinerea* infection of grape berries is associated with activation of lipid metabolism, LOX-related pathway and jasmonate signaling (Agudelo-Romero et al, 2015; Kelloniemi et al, 2015). In grapevine, Girault et al (2008) demonstrated that a lipid transfer protein-JA complex strongly enhanced protection against *B. cinerea*, providing direct evidence that lipid-derived jasmonic acid signaling can contribute to grapevine defense.

Module J was also enriched in cutin biosynthesis, suggesting that infection-associated lipid reprogramming extends to the plant surface. Cuticular lipids are not merely passive physical barriers. Changes in cutin composition and cutin-derived molecules can influence both fungal development and plant immune signaling (Serrano et al, 2014). Importantly, cutin breakdown products can themselves function as damage signals. Cutin monomers have long been proposed as damage-associated molecular patterns. Moreira et al (2024) recently showed that cutin-derived oligomers trigger hallmark immune responses, including Ca^2+^ influx, MAPK activation, ROS production and transcriptional reprogramming characteristic of pattern-triggered immunity. Consistent with this signaling function, experimental alteration of the *Arabidopsis* cuticle, including expression of *B. cinerea* cutinase A, can induce a highly effective resistance state against *B. cinerea*, demonstrating that cuticular perturbation can itself be perceived by the host and activate defense (Chassot et al, 2007). Cutin remodeling during infection may therefore simultaneously affect surface integrity and generate signals reporting damage to the epidermal barrier.

In grapevine, Chavonet et al (2025) recently associated heat-induced tolerance to *B. cinerea* in Cabernet Sauvignon berries with increased cutin monomer accumulation and altered cutin composition. In our system, where reduced symptom development in *V. riparia* was not associated with reduced fungal biomass, induction of cutin-related pathway may therefore contribute to maintenance or restoration of tissue integrity rather than simply preventing fungal penetration. Available comparative anatomy provides only limited support for a constitutive structural explanation. Overall, leaf tissue organization was similar between *V. riparia* and *V. vinifera*, although *V. riparia* displayed a thicker abaxial epidermis in some leaf regions (Paprocka et al, 2023). Comparative microscopy also did not reveal a greater abundance of epicuticular wax in *V. riparia*: dispersed abaxial wax deposit were in fact sparse or absent in both *V. riparia* and *V. vinifera* ‘Cabernet Sauvignon’ (Boso et al, 2010). Nevertheless, biochemical comparisons of whole leaves indicate that wild grapevine, including *V. riparia*, differ quantitatively from domesticated *V. vinifera* in sterol and pentacyclic triterpenoid profiles, compounds that can contribute to cuticular waxes (Burdziej et al, 2019). Because these analyses were performed on whole-leaf extracts rather than isolated cuticles, they provide only indirect evidence that surface lipid composition differs between species. Whether such compositional or epidermal differences contribute to *B. cinerea* penetration, damage perception or cuticular signaling therefore remains unresolved. Lelas et al (2024) further showed that *B. cinerea* degrades plant plasma-membrane glycosylinositol phosphorylceramides through a fungal phospholipase C, illustrating that host lipid compartments themselves are actively remodeled during infection. Together, these findings support a dual role for lipid remodeling during *B. cinerea* infection, coupling oxylipin/jasmonic acid-dependent signaling with dynamic modification of the cuticular and membrane environment.

### 4.5. Cell wall remodeling is a major component of the grapevine response to *B. cinerea*

The cell-wall signatures observed in module H and J suggest that both pre-existing cell-wall properties and infection-induced remodeling may influence the grapevine-*B. cinerea* interaction. Module H associated the basal state of *V. riparia* with primary cell-wall biogenesis and cellulose synthase activity. Cell-wall composition can strongly affect susceptibility to necrotrophs because it determines both the accessibility of polysaccharides to fungal cell-wall degrading enzymes and the production of damage-associated molecular patterns generated during cell wall degradation (Bacete et al, 2018; Rui and Dinneny, 2020). Thus, the constitutive cell-wall-related differences observed in *V. riparia* could influence both tissue resistance to maceration and the perception of plant-induced damage.

During infection, module J revealed a second layer of cell-wall regulation characterized by strong enrichment in pectinesterase and pectinesterase inhibitor activities. This is particularly relevant for *B. cinerea*, whose infection strategy relies heavily on degradation of pectin-rich cell wall matrix. Pectin methylesterification modulates pectin accessibility, mechanical properties and the generation of oligogalacturonide signals, making its dynamic regulation an integral component of cell-wall immunity (Lionetti et al, 2017; Riseh et al, 2024). Functional evidence predating these recent studies already showed that increasing pectin methylesterification through overexpression of pectin methylesterase inhibitors restricts *B. cinerea* growth and disease development in *Arabidopsis* (Lionetti et al, 2007). Conversely, the fungal side of this process is also required for full virulence. Disruption of the *B. cinerea BcPME1* gene reduced total fungal pectin methylesterase activity by approximately 75% and decreased virulence on grapevine as well as on other hosts (Valette-Collet et al, 2003).

Functional evidence in grapevine now directly supports this mechanism. Lagrèze et al (2025) showed that PME10 is strongly induced by *B. cinerea*, that *pme10* reduces pectinesterase activity and increases susceptibility, and that its overexpression reduces disease symptoms. Notably, *VvPME10* is part of our infection-induced module J, and its directly characterized transcriptional regulator *VvWRKY03* is also induced in the three genotypes. Thus, the cell-wall signature detected here includes a regulatory axis with experimentally demonstrated protective activity, rather than only genes annotated for pectin metabolism. Davière et al (2025) further showed that the outcome of the interaction depends not only on host pectin remodeling but also on the structure of the pectin fragments generated by the pathogen. Deletion of the major *B. cinerea* pectin lyase gene *BcPNL1* reduced fungal pathogenicity and delayed both fungal development and activation of host responses. Moreover, the production of pectin-lyase-derived oligogalacturonides was linked to increased expression of JAZ and JOX repressors of jasmonate signaling, thereby connecting pectin degradation directly to manipulation of host hormonal defense. The susceptibility phenotype also depends on pectin esterification properties, emphasizing that the pectin chemistry can determine both fungal degradability and the signaling activity of the resulting fragments. The simultaneous enrichment in pectinesterase and pectinesterase inhibitor activities in module J therefore likely reflect active regulation of pectin methylesterification rather than unidirectional cell wall degradation. These results indicate that pectin remodeling is not a passive consequence of tissue degradation but an active component of grapevine immunity, integrating cell-wall reinforcement, damage-associated molecular patterns generation and modulation of defense signaling.

### 4.6. A distinct immune program in *Vitis riparia* highlights candidate defense-associated genes potentially contributing to tolerance

Unlike the largely conserved response represented by module J, module N displays an infection responsive program preferentially activated in *V. riparia*. This module combines defense, detoxification, secondary metabolism, callose deposition and hormone-related processes and contains several genes annotated as NB-ARC-, NLR-, TIR-NBS-LRR- or RGA-like proteins. Their preferential induction in *V. riparia* suggests that this wild genotype may activate an additional defense-associated transcriptional repertoire potentially contributing to tolerance that is not engaged to the same extent in both *V. vinifera* cultivars.

This hypothesis is consistent with comparative genomics of the genus *Vitis*. Wild grapevines generally carry greater diversity of disease-resistance genes than the cultivated germplasm, and population-scale genome comparisons have shown both a higher abundance and greater diversity contribution of NLR genes in wild grapevine (Long et al, 2024). Comparative genome analysis further revealed major expansion of NLR families in North American wild *Vitis*, including recently duplicated TIR-NLR genes in *V. riparia*, supporting the existence of a particularly diverse receptor repertoire in wild germplasm (Li and Gschwend, 2023).

Evidence specifically related to gray mold also support the contribution of genetically distinct immune programs in wild-derived grapevines. In the highly resistant cultivar Beta, Su et al (2023) identified two stable QTLs for *B. cinerea* resistance on linkage group 2 and 7. Integration with transcriptomics highlighted *VlEDR2*, whose expression was higher in susceptible genotypes, together with candidate regulators *VlERF039*, *VlNAC047* and *VlWRKY51*. Thus, gray mold resistance derived partly from *V. riparia* ancestry has already been genetically associated with a multigenic regulatory architecture rather than a single classical R gene.

A complementary mechanism was identified in resistant *V. amurensis*. Wan et al (2021) found resistance-associated networks containing R proteins, receptor-like kinases, ankyrin proteins and hormone/redox regulators, and functionally demonstrated that overexpression of *VaWRKY10* reduced *B. cinerea* lesions in both *A. thaliana* and susceptible *V. vinifera* ‘Thompson Seedless’. These studies are consistent with module N combining R-gene-like receptors with hormone signaling, detoxification and structural defenses rather than representing an isolated NLR response.

Importantly, resistance to broad-host-range necrotrophs such as *B. cinerea* is generally quantitative and involves many interacting defense processes rather than a simple gene-for-gene relationship. NLR and other immune receptors can contribute to this architecture without necessarily conferring classical monogenic resistance (Corwin et al, 2016). Module N may therefore represent an additional *V. riparia*-specific layer superimposed on the conserved response, combining immune recognition with callose-associated functions, detoxification and jasmonic acid/ethylene-associated defenses. A possible link with symptom limitation comes from a putative *RBOH*, induced at both infection time points specifically in *V. riparia*. This candidate is relevant not simply because RBOHs produce ROS, but because the role of these enzymes in necrotrophic infection can involve redox homeostasis and tissue damage. In *A. thaliana*, disruption of *AtRBOHD* modestly increased susceptibility to *B. cinerea* and impaired the host redox response to its secretome, whereas the fungus could effectively counter extracellular ROS (Vielba-Fernandez et al, 2026). Although the functional relationship between this grapevine locus and *AtDBOHD* remains unverified, these findings provide a plausible framework for investigating whether the additional *V. riparia* response contributes to damage containment without necessarily restricting fungal biomass. Taken together, stronger activation of module N is compatible with a contribution to the progression of symptoms, but does not yet establish a causal tolerance mechanism.

## 5. CONCLUSION

Our results show that contrasting susceptibility to *B. cinerea* among grapevine genotypes cannot be explained simply by differences in the magnitude of a common defense response. The reduced lesion development of *V. riparia*, despite fungal colonization comparable to that of *V. vinifera* at 24 and 48 hpi, was associated with a distinct basal molecular state and an early genotype-specific response, supporting an important disease-tolerance component. At later stages, the three genotypes converged towards a conserved response characterized by repression of photosynthesis- and chloroplast-associated transcriptional programs and coordinated activation of phenylpropanoid/stilbenoid metabolism, lipid-derived signaling and cell-wall remodeling. In addition, *V. riparia* displayed a stronger activation of an immune-associated transcriptional program enriched in defense- and resistance-associated genes potentially contributing to tolerance. Together, these findings indicate that disease outcome is shaped by the temporal organization of defense, from pre-existing molecular states to early genotype-specific responses and conserved late immunity, rather than by response intensity alone. Beyond identifying defense-associated pathways, the transcriptional programs resolved here provide candidate gene sets for functional validation of processes associated with grapevine susceptibility, including potential susceptibility genes whose activity could be targeted or modulated using genome-editing approaches. Recent identification of *VvLYK6* as a negative regulator of chitin-triggered immunity and a candidate susceptibility gene illustrates the potential of this strategy for *B. cinerea* resistance (Villette et al, 2025). Functional validation of candidates emerging from these genotype-dependent programs could ultimately support cultivar-tailored modulation of gray mold susceptibility while limiting disruption of other agronomic traits.

## AUTHOR CONTRIBUTIONS

**Clément Cuello**: Conceptualization, Formal analysis, Data curation, Writing. **Benoit Monnereau**: Investigation, Methodology, Resources. **Samuel Bastos Serra Trinca**: Investigation, Formal analysis. **Cécile Gaillard**: Investigation, Resources. **Vincent Lebeurre**: Investigation, Resources. **Allegra Dovere**: Investigation. **Bouziane Moumen**: Supervision. **Claudia Rouveyrol**: Formal analysis, Investigation, Resources. **Pierre Pétriacq**: Formal analysis, Investigation, Resources. **Pierre Coutos-Thevenot**: Supervision. **Sylvain La Camera**: Conceptualization, Validation, Writing, Funding acquisition.

## ACKNOWLEDGEMENTS

This work was supported by the Centre National de la Recherche Scientifique, the University of Poitiers, the Region Nouvelle-Aquitaine (BM PhD grant - CryptoCARBO project-AAPR2021-2020-11949210), France Relance program, the State-Region Planning Contracts (CPER) and the European Regional Development Fund (FEDER). We are grateful to colleagues for inspiring discussions. We would like to acknowledge the Ebioinfo platform at the University of Poitiers, UMR CNRS 7267, for providing the computational infrastructure and resources essential for the analysis of our data.

## SUPPLEMENTARY MATERIAL

**Supplementary Table S1.** List of primers used for qPCR analysis

**Supplementary Table S2.** Differentially expressed grapevine genes across basal genotype and *Botrytis cinerea* infection contrasts and their SOM module membership

**Supplementary Table S3.** Gene Ontology enrichment of selected SOM transcriptional modules

**Supplementary Table S4.** Correlations between SOM module activity and metabolite abundance across the multi-omics datasets

**Supplementary Figure S1.**
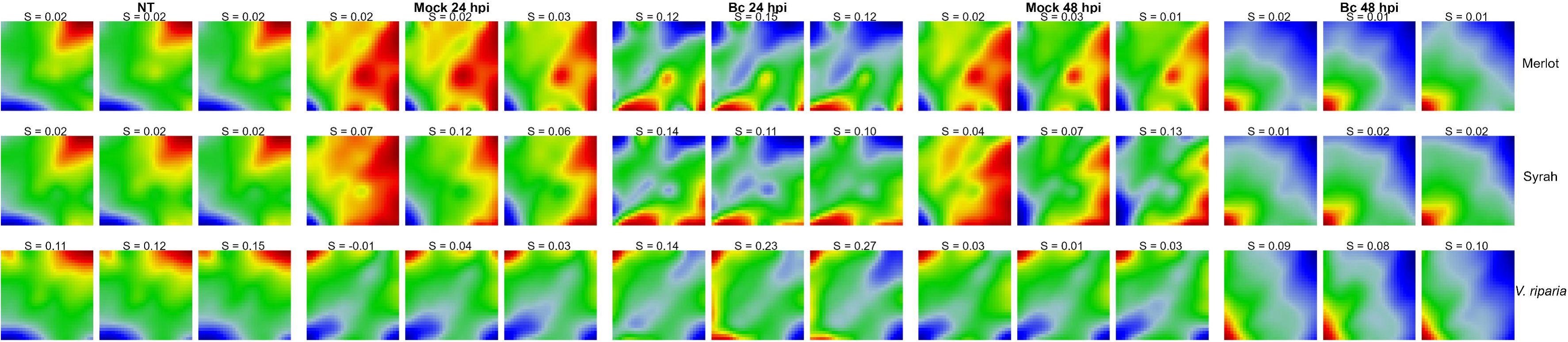
Individual self-organizing map (SOM) expression portraits across grapevine genotypes and experimental conditions. SOM expression portraits are shown for the three biological replicates of <u>V. vinifera</u> ‘Merlot’ and ‘Syrah’ and <u>V. riparia</u> under non-treated conditions (NT), mock treatment at 24 and 48 hpi and <u>B. cinerea</u> infection at 24 and 48 hpi Colors represent relative metagene expression across the SOM landscape, from low expression (blue) to high expression (red). The silhouette score (S) shown above each portrait quantifies the similarity of each sample to its assigned genotype x condition group, with higher values indicating a more distinct and internally consistent expression profile.

**Supplementary Figure S2.**
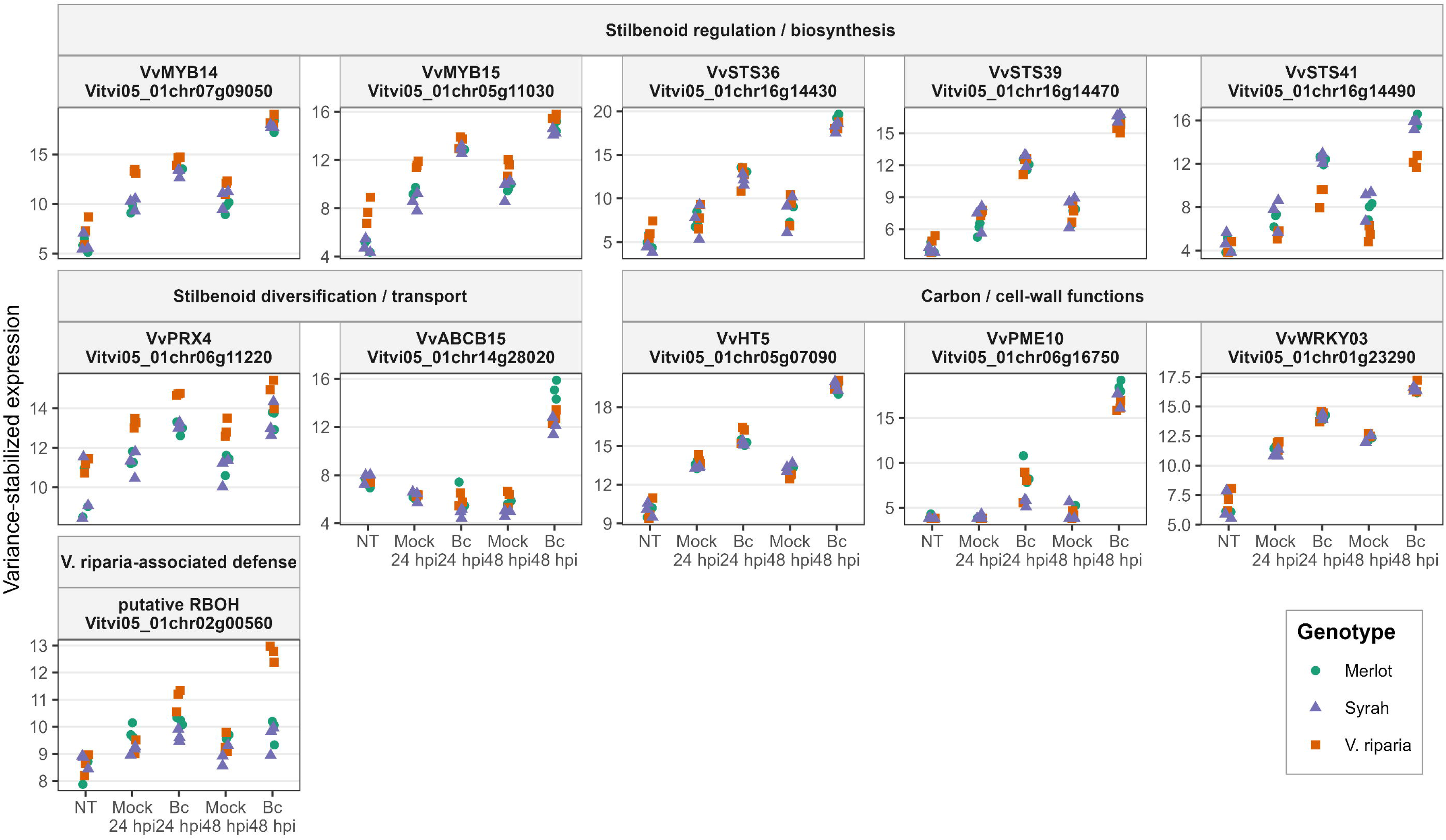
Expression profiles of selected functionally relevant grapevine genes across genotypes and experimental conditions. Genes are grouped according to their functional relevance into stilbenoid regulation/biosynthesis, stilbenoid diversification/transport, carbon/cell-wall functions, and <u>V. riparia</u>-associated defense. Expression values correspond to the three biological replicates of <u>V. vinifera</u> ‘Merlot’, <u>V. vinifera</u> ‘Syrah’ and <u>V. riparia</u> under non-treated conditions (NT), mock treatment at 24 and 48 hpi, and B. cinerea infection (Bc) at 24 and 48 hpi. Each point represents the variance-stabilized expression value of one biological replicate. Colors and symbols indicate grapevine genotype. Gene names and corresponding Vitis vinifera T2T gene identifiers are indicated above each panel.

## REFERENCES

Adrian, M., Jeandet, P., Veneau, J., Weston, L. A., Bessis, R. (1997). Biological activity of resveratrol, a stilbenic compound from grapevines, against Botrytis cinerea, the causal agent for gray mold. Journal of chemical ecology, 23(7), 1689–1702.

Adrian, M., Corio-Costet, M. F., Calonnec, A., Cluzet, S., Poinssot, B., Trouvelot, S., Wiedemann-Merdinoglu, S., Viaud, M. (2024). Grapevine defence mechanisms when challenged by pathogenic fungi and oomycetes. In Advances in botanical research (Vol. 110, pp. 101-195). Academic Press.

Agudelo-Romero, P., Erban, A., Rego, C., Carbonell-Bejerano, P., Nascimento, T., Sousa, L., Martínez-Zapater, J. M., Kopka, J., Fortes, A. M. (2015). Transcriptome and metabolome reprogramming in *Vitis vinifera* cv. Trincadeira berries upon infection with *Botrytis cinerea*. Journal of Experimental Botany, 66(7), 1769–1785.

Alexa, A., Rahnenfuhrer, J., others, 2022. topGO: enrichment analysis for gene ontology. R Package Version 2480 2.

Bacete, L., Mélida, H., Miedes, E., Molina, A. (2018). Plant cell wall□mediated immunity: cell wall changes trigger disease resistance responses. The Plant Journal, 93(4), 614–636.

Bi, K., Liang, Y., Mengiste, T., Sharon, A. (2023). Killing softly: a roadmap of Botrytis cinerea pathogenicity. Trends in plant science, 28(2), 211–222.

Bilgin, D. D., Zavala, J. A., Zhu, J. I. N., Clough, S. J., Ort, D. R., DeLUCIA, E. H. (2010). Biotic stress globally downregulates photosynthesis genes. Plant, cell & environment, 33(10), 1597–1613.

Boso Alonso, S., Alonso-Villaverde Iglesias, V., Santiago Blanco, J. L., Gago Montaña, P., Dürrenberger, M., Düggelin, M., Kassenmeyer, H. H., Martínez Rodríguez, M. D. C. (2010). Macro-and microscopic leaf characteristics of six grapevine genotypes (Vitis spp.) with different susceptibilities to grapevine downy mildew.

Burdziej, A., Pączkowski, C., Destrac-Irvine, A., Richard, T., Cluzet, S., Szakiel, A. (2019). Triterpenoid profiles of the leaves of wild and domesticated grapevines. Phytochemistry Letters, 30, 302–308.

Chassot, C., Nawrath, C., Métraux, J. P. (2007). Cuticular defects lead to full immunity to a major plant pathogen. The Plant Journal, 49(6), 972–980.

Chavonet, E., Mirande-Ney, C., Bernardo, S., Guinand, C., Delestre, G., Zhan, X., Valls Fonayet, J., van Delft, P., Pascal, S., Joubès, J., Lecourieux, D., Fermaud, M., Domergue, F. (2025). Deciphering heat-enhanced resistance of grapevine berries to Botrytis cinerea highlights differential cuticular and secondary metabolite accumulations between Merlot and Cabernet Sauvignon. Plant stress, 101051.

Cheaib, A., Killiny, N. (2025). Photosynthesis responses to the infection with plant pathogens. Molecular Plant-Microbe Interactions, 38(1), 9–29.

Corwin, J. A., Copeland, D., Feusier, J., Subedy, A., Eshbaugh, R., Palmer, C., Maloof, J., Kliebenstein, D. J. (2016). The quantitative basis of the *Arabidopsis* innate immune system to endemic pathogens depends on pathogen genetics. PLoS genetics, 12(2), e1005789.

Coutos□Thévenot, P., Poinssot, B., Bonomelli, A., Yean, H., Breda, C., Buffard, D., Esnault, R., Hain, R., Boulay, M. (2001). In vitro tolerance to Botrytis cinerea of grapevine 41B rootstock in transgenic plants expressing the stilbene synthase Vst 1 gene under the control of a pathogen□inducible PR 10 promoter. Journal of experimental botany, 52(358), 901–910.

Davière, A., Voxeur, A., Jolivet, S., Lelas, L., Vernhettes, S., Soulié, M. C., Fagard, M. (2025). Unravelling the interplay of nitrogen nutrition and the *Botrytis cinerea* pectin lyase BcPNL1 in modulating Arabidopsis thaliana susceptibility. Communications biology, 8(1), 204.

Dubrovina, A. S., Kiselev, K. V. (2017). Regulation of stilbene biosynthesis in plants. Planta, 246(4), 597–623.

Eibach, R., Töpfer, R. (2015). Traditional grapevine breeding techniques. In Grapevine breeding programs for the wine industry (pp. 3-22). Woodhead Publishing.

Gabler, F. M., Smilanick, J. L., Mansour, M., Ramming, D. W., Mackey, B. E. (2003). Correlations of morphological, anatomical, and chemical features of grape berries with resistance to *Botrytis cinerea*. Phytopathology, 93(10), 1263–1273.

Gindro, K., Schnee, S., Righi, D., Marcourt, L., Nejad Ebrahimi, S., Codina, J. M., Voinesco, F., Michellod, E., Wolfender, J. L., Queiroz, E. F. (2017). Generation of antifungal stilbenes using the enzymatic secretome of Botrytis cinerea. Journal of Natural products, 80(4), 887–898.

Girault, T., François, J., Rogniaux, H., Pascal, S., Delrot, S., Coutos-Thévenot, P., Gomès, E. (2008). Exogenous application of a lipid transfer protein-jasmonic acid complex induces protection of grapevine towards infection by *Botrytis cinerea*. Plant Physiology and Biochemistry, 46(2), 140–149.

Haile, Z. M., Pilati, S., Sonego, P., Malacarne, G., Vrhovsek, U., Engelen, K., Tudzynski, P., Zottini, M., Baraldi, E., Moser, C. (2017). Molecular analysis of the early interaction between the grapevine flower and *Botrytis cinerea* reveals that prompt activation of specific host pathways leads to fungus quiescence. Plant, Cell & Environment, 40(8), 1409–1428.

Höll, J., Vannozzi, A., Czemmel, S., D’Onofrio, C., Walker, A. R., Rausch, T., Lucchin, M., Boss, P. K., Dry, I. B., Bogs, J. (2013). The R2R3-MYB transcription factors MYB14 and MYB15 regulate stilbene biosynthesis in Vitis vinifera. The Plant Cell, 25(10), 4135–4149.

Hocini, F. I., Prigent, S., Lerisson, A., Cordazzo, R., Muller, C., Rouveyrol, C., Cassan, C., Rey, A., Gibon, Y., Fuzzati, N., Cocandeau, V., Pétriacq, P. (2026). Metabolomics-based predictive modelling links leaf metabolic profiles to floral traits in Camellia. Journal of Experimental Botany, erag381.

Kelloniemi, J., Trouvelot, S., Héloir, M. C., Simon, A., Dalmais, B., Frettinger, P., Cimerman, A., Fermaud, M., Roudet, J., Baulande, S., Bruel, C., Choquer, M., Couvelard, L., Duthieuw, M., Ferrani, A., Flors, V., Le Pêcheur, P., Loisel, E., Morgant, G., Poussereau, N., Pradier, J. M., Rascle, L., Trdà, L., Poinssot, B., Viaud, M. (2015). Analysis of the molecular dialogue between gray mold (Botrytis cinerea) and grapevine (Vitis vinifera) reveals a clear shift in defense mechanisms during berry ripening. Molecular Plant-Microbe Interactions, 28(11), 1167–1180.

Konecny, T., Nikoghosyan, M., Binder, H. (2023). Machine learning extracts marks of thiamine’s role in cold acclimation in the transcriptome of Vitis vinifera. Frontiers in Plant Science, 14, 1303542.

Konecny, T., Asatryan, A., Nikoghosyan, M., Binder, H. (2024). Unveiling iso-and aniso-hydric disparities in grapevine—a reanalysis by transcriptome portrayal machine learning. Plants, 13(17), 2501.

Konecny, T., Asatryan, A., Binder, H. (2026). Annual transcriptomic trajectory of grapevine buds reveals the coordinated regulation of thiamine and stilbenoid biosynthesis during winter dormancy. BMC Plant Biology, 26(1), 310.

La Camera, S., Gouzerh, G., Dhondt, S., Hoffmann, L., Fritig, B., Legrand, M., & Heitz, T. (2004). Metabolic reprogramming in plant innate immunity: the contributions of phenylpropanoid and oxylipin pathways. Immunological reviews, 198(1), 267–284.

Lagrèze, J., Pajuelo, A. S., Coculo, D., Rojas, B., Pizzio, G. A., Zhang, C., Tian, M. B., Malnoy, M., Vannozzi, A., Dalla Costa, L., Lionetti, V., Tomas Matus, J., Malacarne, G. (2025). PME10 is a pectin methylesterase driving PME activity and immunity against *Botrytis cinerea* in grapevine (*Vitis vinifera* L.). Plant Biotechnology Journal, 23(11), 4981–4997.

Langcake, P. (1981). Disease resistance of *Vitis* spp. and the production of the stress metabolites resveratrol, ε-viniferin, α-viniferin and pterostilbene. Physiological Plant Pathology, 18(2), 213–226.

Le Roy, J., Huss, B., Creach, A., Hawkins, S., Neutelings, G. (2016). Glycosylation is a major regulator of phenylpropanoid availability and biological activity in plants. Frontiers in plant science, 7, 735.

Lelas, L., Rouffet, J., Filachet, A., Sechet, J., Davière, A., Desprez, T., Vernhettes, S., Voxeur, A. (2024). A fungal phospholipase C involved in the degradation of plant glycosylinositol phosphorylceramides during *Arabidopsis*/*Botrytis* interaction. Communications Biology, 7(1), 1372.

Li, B., Gschwend, A. R. (2023). *Vitis labrusca* genome assembly reveals diversification between wild and cultivated grapevine genomes. Frontiers in Plant Science, 14, 1234130.

Lionetti, V., Raiola, A., Camardella, L., Giovane, A., Obel, N., Pauly, M., Favaron, F., Cervone, F., Bellincampi, D. (2007). Overexpression of pectin methylesterase inhibitors in Arabidopsis restricts fungal infection by Botrytis cinerea. Plant physiology, 143(4), 1871–1880.

Lionetti, V., Fabri, E., De Caroli, M., Hansen, A. R., Willats, W. G., Piro, G., Bellincampi, D. (2017). Three pectin methylesterase inhibitors protect cell wall integrity for *Arabidopsis* immunity to *Botrytis*. Plant Physiology, 173(3), 1844–1863.

Löffler-Wirth, H., Kalcher, M., Binder, H. (2015). oposSOM: R-package for high-dimensional portraying of genome-wide expression landscapes on bioconductor. Bioinformatics, 31(19), 3225–3227.

Long, Q., Cao, S., Huang, G., Wang, X., Liu, Z., Liu, W., Wang, Y., Xiao, H., Peng, Y., Zhou, Y. (2024). Population comparative genomics discovers gene gain and loss during grapevine domestication. Plant Physiology, 195(2), 1401–1413.

Love, M. I., Huber, W., Anders, S. (2014). Moderated estimation of fold change and dispersion for RNA-seq data with DESeq2. Genome biology, 15(12), 550.

Maggiolini, F. A. M., Prencipe, A., Bergamini, C., Marsico, A. D., Vendemia, M., Santamaria, M., Giannandrea, M. A., D’Amico, M., Forleo, L. R., Perniola, R., Velasco, R., Cardone, M. F. (2025). A comparative transcriptomic study reveals temporal and genotype-specific defense responses to *Botrytis cinerea* in Grapevine. Journal of Fungi, 11(2), 124.

Maia, M., Aziz, A., Jeandet, P., Carré, V. (2023). Profiling and localization of stilbene phytoalexins revealed by MALDI-MSI during the grapevine–*Botrytis cinerea* interaction. Journal of Agricultural and Food Chemistry, 71(42), 15569–15581.

Marant, B., Flourat, A. L., Lanoue, A., Magot, F., Parent, L., Leclère, V., Guihard, G., Aggad, D., Jacquard, C., Courot, E., Aziz, A., Crouzet, J. (2025). Peroxidase 4-based enzymatic synthesis of stilbene oligomers in methyl jasmonate-elicited grapevine cell suspensions. Journal of Agricultural and Food Chemistry, 73(3), 1929–1939.

Martínez-Márquez, A., Martins, V., Sellés-Marchart, S., Gerós, H., Corchete, P., Bru-Martínez, R. (2025). The grapevine ABC transporter B family member 15 is a trans-resveratrol transporter out of grapevine cells. Frontiers in Plant Science, 15, 1450638.

Monnereau, B., Cuello, C., Gaillard, C., Lebeurre, V., Videau, P., Zekri, O., Coutos-Thévenot, P., La Camera, S. (2026). Overexpression of the sugar transporter VvHT5 in grapevine drives coordinated host–pathogen transcriptional reprogramming favoring *Botrytis cinerea*. Plant Physiology and Biochemistry, 111480.

Monnereau, B., Cuello, C., Bastos Serra Trinca, S., Gaillard, C., Lebeurre, V., Dovere, A., Moumen, B., Rouveyrol, C., Pétriacq, P., Coutos-Thevenot, P., La Camera, S. Transcriptome and metabolome datasets of *Botrytis cinerea*-infected grapevine leaves from *Vitis vinifera* ‘Merlot’ and ‘Syrah’, and *Vitis riparia* ‘Gloire de Montpellier’. Data in Brief, submitted

Morant, A. V., Jørgensen, K., Jørgensen, C., Paquette, S. M., Sánchez-Pérez, R., Møller, B. L., Bak, S. (2008). β-Glucosidases as detonators of plant chemical defense. Phytochemistry, 69(9), 1795–1813.

Moreira, C. J., Escórcio, R., Bento, A., Bjornson, M., Herold, L., Tomé, A. S., Martins, C., Fanuel, M., Matrins, I., Bakan, B., Zipfel, C., Silva Pereira, C. (2024). Cutin-derived oligomers induce hallmark plant immune responses. Journal of experimental botany, 75(16), 5146–5161.

Pagán, I., García-Arenal, F. (2020). Tolerance of plants to pathogens: a unifying view. Annual review of phytopathology, 58(1), 77–96.

Paprocka, M., Dancewicz, K., Kordan, B., Damszel, M., Sergiel, I., Biesaga, M., Mroczek, J., Arroyo Garcia, R. A., Gabryś, B. (2023). Probing behavior of Aphis fabae and Myzus persicae on three species of grapevines with analysis of grapevine leaf anatomy and allelochemicals. The European Zoological Journal, 90(1), 83–100.

Pedras, M. S. C., Yaya, E. E. (2015). Plant chemical defenses: are all constitutive antimicrobial metabolites phytoanticipins?. Natural Product Communications, 10(1), 1934578X1501000142.

Pezet, R., Perret, C., Jean-Denis, J. B., Tabacchi, R., Gindro, K., Viret, O. (2003a). δ-Viniferin, a resveratrol dehydrodimer: one of the major stilbenes synthesized by stressed grapevine leaves. Journal of agricultural and Food Chemistry, 51(18), 5488–5492.

Pezet, R., Viret, O., Perret, C., Tabacchi, R. (2003b). Latency of Botrytis cinerea Pers.: Fr. and biochemical studies during growth and ripening of two grape berry cultivars, respectively susceptible and resistant to grey mould. Journal of Phytopathology, 151(4), 208–214.

Piasecka, A., Jedrzejczak□Rey, N., Bednarek, P. (2015). Secondary metabolites in plant innate immunity: conserved function of divergent chemicals. New Phytologist, 206(3), 948–964.

Polesani, M., Bortesi, L., Ferrarini, A., Zamboni, A., Fasoli, M., Zadra, C., Lovato, A., Pezzotti, M., Delledonne, M., Polverari, A. (2010). General and species-specific transcriptional responses to downy mildew infection in a susceptible (*Vitis vinifera*) and a resistant (*V. riparia*) grapevine species. BMC genomics, 11(1), 117.

Rahman, M. U., Ma, Q., Ahmad, B., Hanif, M., Zhang, Y. (2020). Histochemical and microscopic studies predict that grapevine genotype “ju mei gui” is highly resistant against *Botrytis cinerea*. Pathogens, 9(4), 253.

Rahman, M. U., Liu, X., Wang, X., Fan, B. (2024). Grapevine gray mold disease: infection, defense and management. Horticulture Research, 11(9), uhae182.

Riseh, R. S., Vazvani, M. G., Taheri, A., Kennedy, J. F. (2024). Pectin-associated immune responses in plant-microbe interactions: A review. International Journal of Biological Macromolecules, 273, 132790.

Rossi, F. R., Krapp, A. R., Bisaro, F., Maiale, S. J., Pieckenstain, F. L., Carrillo, N. (2017). Reactive oxygen species generated in chloroplasts contribute to tobacco leaf infection by the necrotrophic fungus *Botrytis cinerea*. The Plant Journal, 92(5), 761–773.

Rui, Y., Dinneny, J. R. (2020). A wall with integrity: surveillance and maintenance of the plant cell wall under stress. New Phytologist, 225(4), 1428–1439.

Rui, L., Yang, S. Q., Zhou, X. H., Wang, W. (2025). The important role of chloroplasts in plant immunity. Plant Communications, 6(8).

Schneider, C. A., Rasband, W. S., Eliceiri, K. W. (2012). NIH Image to ImageJ: 25 years of image analysis. Nature methods, 9(7), 671–675.

Serrano, M., Coluccia, F., Torres, M., L’Haridon, F., Métraux, J. P. (2014). The cuticle and plant defense to pathogens. Frontiers in plant science, 5, 274.

Soares, F., Pimentel, D., Erban, A., Neves, C., Reis, P., Pereira, M., Rego, C., Gama-Carvalho, M., Kopka, J., Fortes, A. M. (2022). Virulence-related metabolism is activated in *Botrytis cinerea* mostly in the interaction with tolerant green grapes that remain largely unaffected in contrast with susceptible green grapes. Horticulture Research, 9, uhac217.

Staats, M., van Kan, J. A. (2012). Genome update of Botrytis cinerea strains B05. 10 and T4.

Su, K., Zhao, W., Lin, H., Jiang, C., Zhao, Y., Guo, Y. (2023). Candidate gene discovery of *Botrytis cinerea* resistance in grapevine based on QTL mapping and RNA-seq. Frontiers in plant science, 14, 1127206.

Sun, Q., Hou, S., Gao, H. (2025). Chloroplasts orchestrate multilayered defences against plant pathogens. Plant, Cell & Environment, 48(12), 8502–8517.

Taillis, D., Becissa, O., Pébarthé-Courrouilh, A., Renouf, E., Palos-Pinto, A., Richard, T., Cluzet, S. (2023). Antifungal activities of a grapevine byproduct extract enriched in complex stilbenes and stilbenes metabolization by *Botrytis cinerea*. Journal of Agricultural and Food Chemistry, 71(11), 4488–4497.

Umberath, K. M., Mischke, A., Caspers-Weiffenbach, R., Backmann, L., Scharfenberger-Schmeer, M., Wegmann-Herr, P., Schieber, A., Weber, F. (2024). Curse or blessing: Growth-and laccase-modulating properties of polyphenols and their oxidized derivatives on *Botrytis cinerea*. Food Research International, 192, 114782.

Umberath, K. M., Backmann, L., Scharfenberger□Schmeer, M., Wegmann□Herr, P., Schieber, A., Weber, F. (2026). Phenolic profile of selected grape (*Vitis vinifera* L.) varieties in the course of ripening and *Botrytis cinerea* infections. Journal of the Science of Food and Agriculture.

Valette-Collet, O., Cimerman, A., Reignault, P., Levis, C., Boccara, M. (2003). Disruption of Botrytis cinerea pectin methylesterase gene Bcpme1 reduces virulence on several host plants. Molecular Plant-Microbe Interactions, 16(4), 360–367.

Veloso, J., van Kan, J. A. (2018). Many shades of grey in Botrytis–host plant interactions. Trends in plant science, 23(7), 613–622.

Vielba-Fernandez, A., Sommer, L., Ross, D., Ingelfinger, J., Wachtmeister, T., Hecht, J., Safari, N., Müller-Schüssele, S. J., Hahn, M. (2026). Botrytis cinerea uses effective detoxification mechanisms to overcome the plant RBOHD-dependent extracellular oxidative burst. Frontiers in Plant Science, 17, 1969467.

Villette, J., Marzari, T., Landry, D., Roudaire, T., Klinguer, A., Leborgne-Castel, N., Vicedo, C., Gasciolli, V., Pouzet, C., Lefebvre, B., Héloir, M. C., Poinssot, B. (2025). The Vitis vinifera receptor VvLYK6 negatively regulates chitin-triggered immune responses and promotes fungal infections. Frontiers in Plant Science, 16, 1705961.

Wan, R., Hou, X., Wang, X., Qu, J., Singer, S. D., Wang, Y., Wang, X. (2015). Resistance evaluation of Chinese wild *Vitis* genotypes against *Botrytis cinerea* and different responses of resistant and susceptible hosts to the infection. Frontiers in Plant Science, 6, 854.

Wan, R., Guo, C., Hou, X., Zhu, Y., Gao, M., Hu, X., Zhang, S., Jiao, C., Guo, R., Li, Z., Wang, X. (2021). Comparative transcriptomic analysis highlights contrasting levels of resistance of *Vitis vinifera* and *Vitis amurensis* to *Botrytis cinerea*. Horticulture research, 8.

Wan, R., Yang, Z., Liu, J., Zhang, M., Jiao, J., Wang, M., Zhang, K., Hao, P., Liu, Y., Bai, T., Song, C., Song, S., Shi, J., Zheng, X. (2024). Identification of Laccase Genes in Grapevine and Their Roles in Response to Botrytis cinerea. Horticulturae, 10(4), 376.

Williamson, B., Tudzynski, B., Tudzynski, P., Van Kan, J. A. (2007). Botrytis cinerea: the cause of grey mould disease. Molecular plant pathology, 8(5), 561–580.

Windram, O., Madhou, P., McHattie, S., Hill, C., Hickman, R., Cooke, E., Jenkins, D. J., Penfold, C. A., Baxter, L., Breeze, E., Kiddle, S. J., Rhodes, J., Atwell, K., Kliebenstein, D. J., Kim, Y., Stegle, O., Borgwardt, K., Zhang, C., Tabrett, A., Legaie, R., Moore, J., Finkenstadt, B., Wild, D. L., Mead, A., Beynon, J., Ott, S., Buchanan-Wollaston, V., Denby, K. J. (2012). *Arabidopsis* defense against *Botrytis cinerea*: chronology and regulation deciphered by high-resolution temporal transcriptomic analysis. The Plant Cell, 24(9), 3530–3557.

Zhang, W., Wang, S., Yang, J., Kang, C., Huang, L., Guo, L. (2022). Glycosylation of plant secondary metabolites: Regulating from chaos to harmony. Environmental and Experimental Botany, 194, 104703.

Zimdars, S., Hitschler, J., Schieber, A., Weber, F. (2017). Oxidation of wine polyphenols by secretomes of wild *Botrytis cinerea* strains from white and red grape varieties and determination of their specific laccase activity. Journal of Agricultural and Food Chemistry, 65(48), 10582–10590.

